# IL-23 drives uveitis by acting on a novel population of tissue-resident entheseal T cells

**DOI:** 10.1101/2024.05.16.594586

**Authors:** Robert Hedley, Amy Ward, Colin J Chu, Sarah E Coupland, Serafim Kiriakidis, Peter C Taylor, Stephanie G Dakin, The ORBIT consortium, Christopher D Buckley, Jonathan Sherlock, Andrew D Dick, David A Copland

## Abstract

Recurrent acute anterior uveitis is a frequent extra-articular manifestation of the axial spondyloarthropathies (AxSpA); chronic inflammatory diseases affecting the spine, enthesis, peripheral joints, skin, and gastrointestinal tract. Pathology in AxSpA has been associated with local tissue-resident populations of interleukin (IL)-23 responsive lymphoid cells. Here we reveal a novel population of ocular T cells defined by CD3+CD4-CD8-γδTCR+IL-23R+ that reside within the anterior uvea as an ocular entheseal analogue of the mouse eye. Localised cytokine expression demonstrates that uveal IL-23R+ IL-17A-producing cells are both necessary and sufficient to drive uveitis in response to IL-23. This T cell population is also present in humans, occupying extravascular tissues of the anterior uveal compartment. Consistent with the concept of IL-23 as a unifying mediator in AxSpA, we present evidence that IL-23 can also act locally on tissue resident T cells in the anterior compartment of the eye at sites analogous to the enthesis to drive ocular inflammation.

## Introduction

Axial spondyloarthropathies (AxSpA) represent a group of chronic immune-mediated inflammatory diseases sharing clinical and molecular similarities, predominantly involving the axial skeleton but also peripheral joints and entheses (ReA) [1–3]. Frequently associated with these conditions are extra-articular manifestations including inflammation of the skin (psoriasis), gut (Inflammatory bowel disease; IBD) and eye (uveitis) [4, 5]. Further evidence that these conditions are unified at a cellular and mechanistic level comes from the observations that in patients with one disease, subclinical disease is often present at other anatomical sites. For example, AxSpA is strongly associated with subclinical gut inflammation [6] and IBD, and psoriasis are associated with subclinical entheseal inflammation [7].

Although the underlying pathogenic mechanisms of AxSpA are still not fully understood, strong genetic links implicate both human leukocyte antigen B*27 (HLA-B*27) allele, and the interleukin IL-23 pathway [8–17]. The importance of IL-23 as a unifying factor for AxSpA is highlighted by single nucleotide polymorphisms (SNPs) in the IL-23 receptor (IL-23R) and the IL-23 cytokine, in genes involved in downstream signalling pathways and the IL-17 axis [18]. The IL-23 pathway is recognised to play a prominent role at externally facing barrier surfaces, particularly the skin [19] and gut [20] but can also drive inflammation at internal sterile sites such as the joints [21]. Whilst barrier surfaces are characterized by the presence of an extensive microbiome, a fundamental feature of the joints is the presence of high biomechanical stress and tension.

IL-23R is constitutively expressed on various immune cell populations, including natural killer (NK) cells, innate lymphoid cells, γδ T cells, and mucosal-associated invariant T (MAIT) cells, all of which recognize structural elements via invariant T cell receptors or other recognition motifs. Engagement of IL-23R activates the intracellular Janus kinase-signal transducer and activator of transcription (JAK-STAT) signalling pathway with tyrosine-protein kinase (TYK2) and STAT3 being the dominant drivers for pathogenic T helper 17 (Th17) cell cytokines (e.g. IL-17, IL-22, GM-CSF) which promote chronic tissue inflammation [22, 23]. The ability of IL-23 to act rapidly at mucosal barrier tissues is partly due to the presence of resident type 17 cells which express IL-23R [24]. Tissue-resident IL-23 responsive CD3+CD4-CD8-T cells are found at highly defined positions of the musculoskeletal (MSK) entheses in mice [21] and humans [25], and other anchorage points associated with biomechanical stress including the aortic root [26, 27]. Intriguingly, these tissue-resident cells are present even in the healthy state which indicates their potential role in regulating barrier function, tissue repair, and homeostasis. The pattern of tissue localization of IL-23R expressing cells can therefore determine how dysregulation of IL-23 biology can elicit inflammation at these precise anatomical sites.

Uveitis represents a heterogenous group of inflammatory disorders characterized by infiltration of leukocytes into the uveal tissues and intraocular cavities of the eye [28]. Anterior uveitis (AU), primarily affecting the iris and ciliary body, represents the most frequent type (approximately 80% of cases), resulting in vision loss through, for example, secondary to cataracts, glaucoma, or macular oedema [29–31]. Acute anterior uveitis (AAU) is the most severe form [32], presenting with acute onset of discomfort, eye redness, visual impairment, and cellular infiltration in the aqueous humor (AqH). AAU is classically described as the most common extra-articular manifestation in AxSpA, with a third of patients developing intraocular inflammation [33]. Patients with apparently isolated uveitis not only have a tendency for subclinical bowel inflammation [34], but also extensive subclinical enthesitis [35]. HLA-B27-associated uveitis is often undiagnosed and consequently, its association with AxSpA is overlooked [36, 37]. HLA-B27 protein, present in up to 50% of patients with AAU, can misfold triggering the unfolded protein response resulting in the production of IL-23 [38]. In human studies, elevated serum levels of IL-23 are associated with an increased risk of AAU in patients with AxSpA [39] as well as other forms of uveitis including Vogt-Koyanagi-Harada (VKH) and Behçet’s disease (BD) [40–42]. Human genome-wide association studies (GWAS) demonstrate that in patients SNPs in the IL-23R gene are associated with uveitis [43, 44]. Irrespective of HLA-B27 positivity, AAU is also a feature of AxSpA-related diseases.

Collectively, these strong associations suggest common factors responsible for disease, with pathology orchestrated by resident populations of IL-23 responsive cells. Therapies neutralising IL-23 in human patients are effective in psoriasis, PsA, and IBD [45, 46]. However, despite successes and insights, the biology and immunological aetiology of human uveitis has remained enigmatic, in part due to the difficulty of obtaining healthy eye tissue and samples. We, therefore, sought evidence for a resident population of IL-23 responsive cells within the eye analogous to musculoskeletal entheses. We show that IL-23 promotes intraocular inflammation in the mouse eye by acting on a previously unidentified population of CD3+CD4-CD8-γδTCR+IL-23R+ cells resident in the anterior uvea, and that the expression of this cytokine alone, in the absence of other inflammatory signals, is sufficient to reproduce classical features of uveitis. Furthermore, data from post-mortem human tissues demonstrates the extravascular location of resident CD3+γδTCR+ cells in the ciliary body and sclera, which secrete IL-17A upon activation. Our data supports that IL-23/IL-17 axis is an important therapeutic target in uveitis.

## Results

### Tissue-resident IL-23R+ γδ T-cells are located in the murine anterior uvea

Using light sheet fluorescence microscopy (LSFM), we assessed whether T cell populations are present within the anterior uvea and iridocorneal angle tissue in adult albino (B6(Cg)-Tyrc-2J/J) mice. Perfused anterior segments were optically cleared [47], immunolabelled with anti-CD3 antibody and DAPI, before 3D cross-sectional images of the anterior chamber from different angles were captured. Low magnification LSFM imaging of the anterior structure demonstrated clusters of CD3+ T cells located within different regions including the corneal limbus, sclera and ciliary body **(Figure 1A)**.

**Figure 1:**
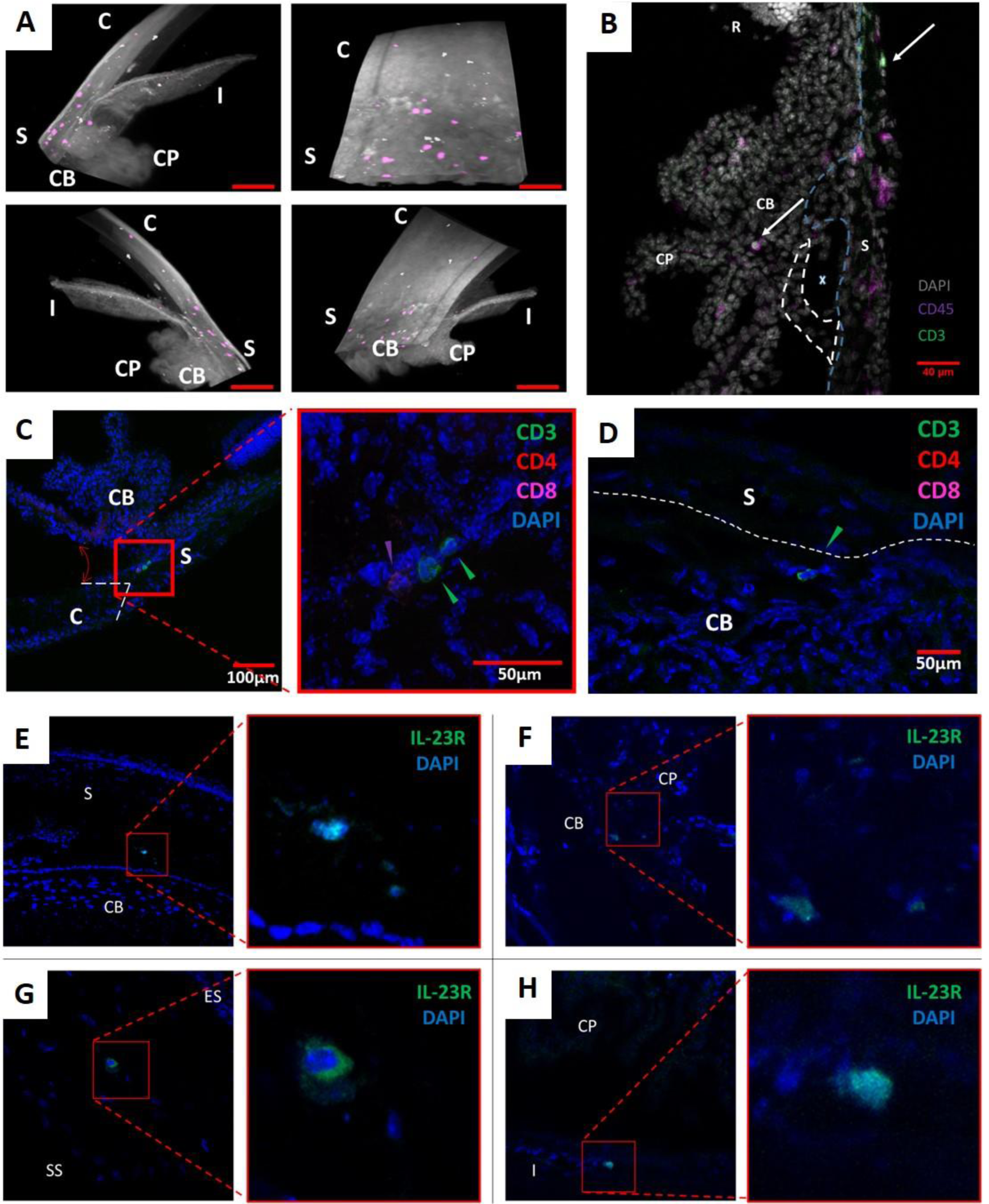
Tissue-resident CD3+IL-23R+ T cells are found in the mouse anterior uvea. Naïve B6(Cg)-Tyrc-2J/J (albino) mice were perfused, and intact globes optically cleared, immunolabelled with CD3e and DAPI. **(A)** Lightsheet Z.1 acquired, immunofluorescent whole mount images, (ii) rotated 90°, (iii) rotated 180°, and (iv) rotated 270°. 3D rendered image shows expression of tissue autofluorescence (white) and CD3e (purple). Co-expression of CD3 and nuclei observed within different regions including the peripheral cornea, limbal sclera and ciliary body. Images captured at x10 magnification. Red scale bar: 100μm. CP – ciliary process, CB – ciliary body, S – sclera, I – iris, and C – cornea. **(B)** Immunofluorescence image of anterior tissue section from perfused albino mouse demonstrates presence of CD45+CD3+ T cells within extravascular tissues including the trabecular meshwork, ciliary body and sclera. [Image captured at x20 magnification. Dashed blue line – the inner limit of the sclera, dashed white line – trabecular meshwork, X – Schlemm’s canal, P – ciliary process, CB – ciliary body, S –sclera, and R – retina]. **(C)** CD4+ and CD8+ (single positive) cells are restricted to the juxtacanalicular tissue (JCT) region of the trabecular meshwork, proximal to the inner wall of Schlemm’s canal. Coloured arrows indicating T cell subtypes; CD3+ (green), CD4 (red), and CD8+ (purple). **(D)** Sections also highlighted CD3+CD4-CD8-(double negative T cells) in the region of ciliary body proximal to the limbal sclera. Sections taken from naïve IL-23R-eGFP(+/-) reporter mice IL-23R+ cells are evident within the limbal sclera, specifically the transitional region of the sclera to ciliary body **(E)** and the transitional region from episclera (ES) to stromal sclera (SS) and not the vascular ES **(F)**, the folding of the inner ciliary body **(G)**, and posterior border layer of the iris **(H)**. Nuclei (blue) and IL-23R-eGFP (green). Red square highlights region of tissue magnified in adjacent image **(C, E-H)**.

To determine their precise tissue location and phenotype, serial sections from perfused eyes were immunostained with additional surface markers, demonstrating that CD45+CD3+ cells reside within extravascular tissues including the trabecular meshwork, ciliary body, and sclera **(Figure 1B)**. CD4+CD8-Th cells and CD4-CD8+ cytotoxic T cell (CD8+) populations are restricted to the juxtacanalicular tissue (JCT) region of the trabecular meshwork (TM), proximal to the inner wall of Schlemm’s canal **(Figure 1C)**. Interestingly, a population of CD3+CD4-CD8-(double negative T cells) were identified in the region of the ciliary body and irido-corneal angle **(Figure 1D)**, present in both the ciliary body and sclera parallel to the *pars plana* and *pars plicata* regions. Furthermore, using IL-23R-eGFP reporter mice, IL-23R+ cells are evident within the limbal sclera **(Figure 1E)** - specifically within the scleral stroma proper and not the vascular episclera **(Figure 1F)** - the ciliary body **(Figure 1G)**, and posterior border of the iris **(Figure 1H)**.

To define the phenotype and frequency of CD3+ populations, naïve IL-23R-eGFP+/- (heterozygous) mouse eyes were prepared for flow cytometry **(Figure 2A)**. Enzymatic digestion of dissected anterior uveal tissue (including the limbal sclera, cornea, iris, and ciliary body) reveals that the IL-23R+ fraction represents 10-15% of the total CD3+ T cell pool, and in absolute numbers 50 cells can be routinely isolated from a single anterior uvea sample **(Figure 2B)**. Furthermore, comparison of heterozygous and homozygous IL-23R-eGFP eyes reveals that IL-23R expression may represent a critical determinant in the residency/accumulation of this T cell population within a specific ocular location. In IL-23R-eGFP+/+ (homozygous) in which eGFP reporter sequences replace both IL-23R coding sequences rendering these mice functionally deficient in IL-23R, the total number and frequency of CD3+IL23R+ T cells is significantly reduced **(Supplementary Figure 1A).**

**Figure 2:**
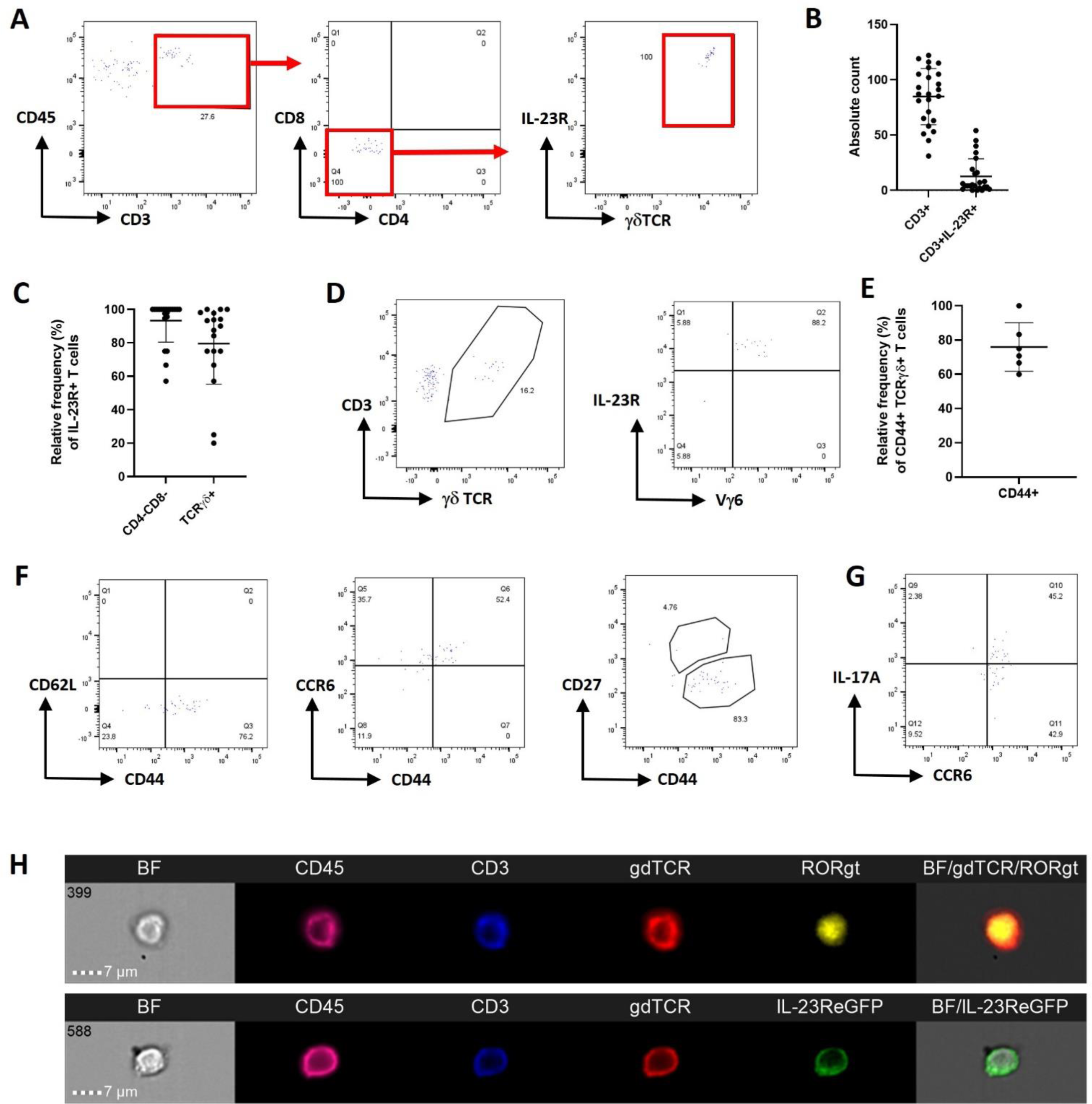
Naïve anterior uvea contains CD3+γδTCR+IL-23R+ T cells with intrinsic pathogenic capacity. Anterior uvea tissue prepared from naïve IL-23R-eGFP (+/-) reporter mice were analysed by flow cytometry. **(A)** Representative flow plots showing gating strategy which identifies resident T cells as CD3+CD4-CD8-IL-23R+γδTCR+. **(B)** Graphs showing absolute counts of CD3+ and CD3+IL-23R+ populations isolated, and **(C)** relative frequency of IL-23R+ cells expressing γδTCR+ (n=25 AU samples). **(D)** Flow plots demonstrating CD3+IL-23R+γδTCR+ cells are Vγ6+ (no detectable expression of Vγ1 or Vγ4 chains), indicating an IL-17+ producing subset. **(E&F)** Resident IL-23R+γδTCR cells display a CD44^high^CD62L-(effector phenotype), expressing CCR6+ but not CD27. **(G)** Following *ex vivo* stimulation (PMA/Ionomycin) of single cell anterior uvea suspensions, intracellular cytokine staining demonstrates cells only secrete IL-17A. **(H)**. Imagestream analysis of pooled naïve anterior uvea suspensions (n=4) confirms CD3+ IL-23R+ γδ+ surface phenotype, and intracellular expression of RORγT.

Consistent with published observations of musculoskeletal (MSK) entheseal tissues [21, 26], CD3+CD4-CD8-γδ T-cells, but not αβ T cells, constitute the majority of anterior compartment IL-23R+ T cells **(Figure 2C)**. Typically, γδ T-cell populations are classified according to the expression of the variable domain of the TCRγ (Vγ) chain [48]. In the anterior compartment, all isolated γδ T cells were Vγ6+, with no detectable expression of Vγ1 or Vγ4 chains **(Figure 2D; Supplementary 1B)**, suggesting an IL-17+ producing subset, as described for the MSK entheseal tissues, reproductive tract, lung, and skin [49, 50].

### Resident IL-23R+ γδ+ T cells exhibit a pre-activated effector phenotype

Expression of IL-23R is a recognised hallmark of IL-17-producing cells, including Th17 cells [51–53], γδ T cells [54], and Type 3 Innate Lymphoid Cells (ILC3) [55, 56]. Furthermore, expression of C-C chemokine receptor 6 (CCR6) by T cells, including IL-17 producing γδ T cells (γδ17) is linked to increased IL-17A secretion [57]. We therefore sought to chacterize the IL-23R+ γδ+ T cell phenotype to define their functional capacity.

Activated and functionally differentiated effector γδ T cells express high levels of CD44 [49]. In naïve mice, ∼95% of anterior compartment-resident γδ T cells are CD44^high^ **(Figure 2E)**, and display a CD44^high^CD62L-effector phenotype **(Figure 2F)**. Expression of CCR6 and CD27 permits functional discrimination between IL-17-producing and IFNγ-producing γδ T cell subsets [58]. In the anterior compartment, the γδ T cells are CCR6+CD27-**(Figure 2F)**, and following *ex vivo* stimulation, secrete IL-17A but not IFN-γ **(Figure 2G; Supplementary Figure 1C)**. Furthermore, ImageStream analysis confirmed the CD3+ IL-23R+ γδ+ surface phenotype, and importantly highlights the intracellular expression/localisation of retinoid orphan receptor gamma t (RORγt) in cells isolated from the naïve anterior compartment **(Figure 2H)**.

These data demonstrate that healthy anterior uvea contains a resident T cell population, defined by surface marker expression, CD3+IL-23R+Vγ6+ γδ+CCR6+. These cells are equipped with an intrinsic capacity to secrete IL-17A, and therefore a potential to act as pathogenic effectors in the eye, comparable to the pathogenic cells described in psoriasis and in the MSK enthesis [21, 26, 59].

### IL-23 overexpression alone in the eye is sufficient to drive inflammation *in vivo*

To evaluate IL-23 responsiveness of this resident T cell population *in vivo*, we engineered a ShH10 serotype adeno-associated virus (AAV) encoding a ‘hyper-IL-23’ cytokine [60] transgene to facilitate localized secretion of cytokine within the mouse eye. The ShH10 capsid serotype permits rapid transduction of the ciliary body non-pigmented epithelium and cells of the inner retina (Müller glia, ganglion cells, and astrocytes) when injected intravitreally into the mouse eye [61].

Following intravitreal injection of 1×10^11^ vector genomes (vg)/eye of ShH10_IL-23 vector in C57BL/6J (wild-type) mice, posterior (retinal) inflammation, subtle optic disc swelling, retinal vascular changes, and cellular infiltration of the vitreous and aqueous cavities are manifest by day 12 **(Figure 3A)**. In eyes receiving a control virus (ShH10_GFP; expressing a widely used reporter protein (GFP) and accepted to be non-inflammatory) or vehicle (PBS) injection, no clinical inflammation or pathological changes are observed at the same time-point. We performed immune profiling using flow cytometry to characterize the immune cell populations infiltrating both the anterior and posterior compartments. This demonstrates that IL-23 overexpression leads to a significant increase of CD45+ cells in each ocular compartment **(Figure 3B)**, comprising both adaptive but also innate immune populations **(Supplementary Figure 2)**. ShH10_GFP (control AAV) also results in subclinical accumulation of CD45+ cells compared to the naïve eye, despite no obvious signs of inflammation, which reflects recognized AAV-mediated changes in the immune threshold of the tissue following intravitreal injection [62, 63].

**Figure 3:**
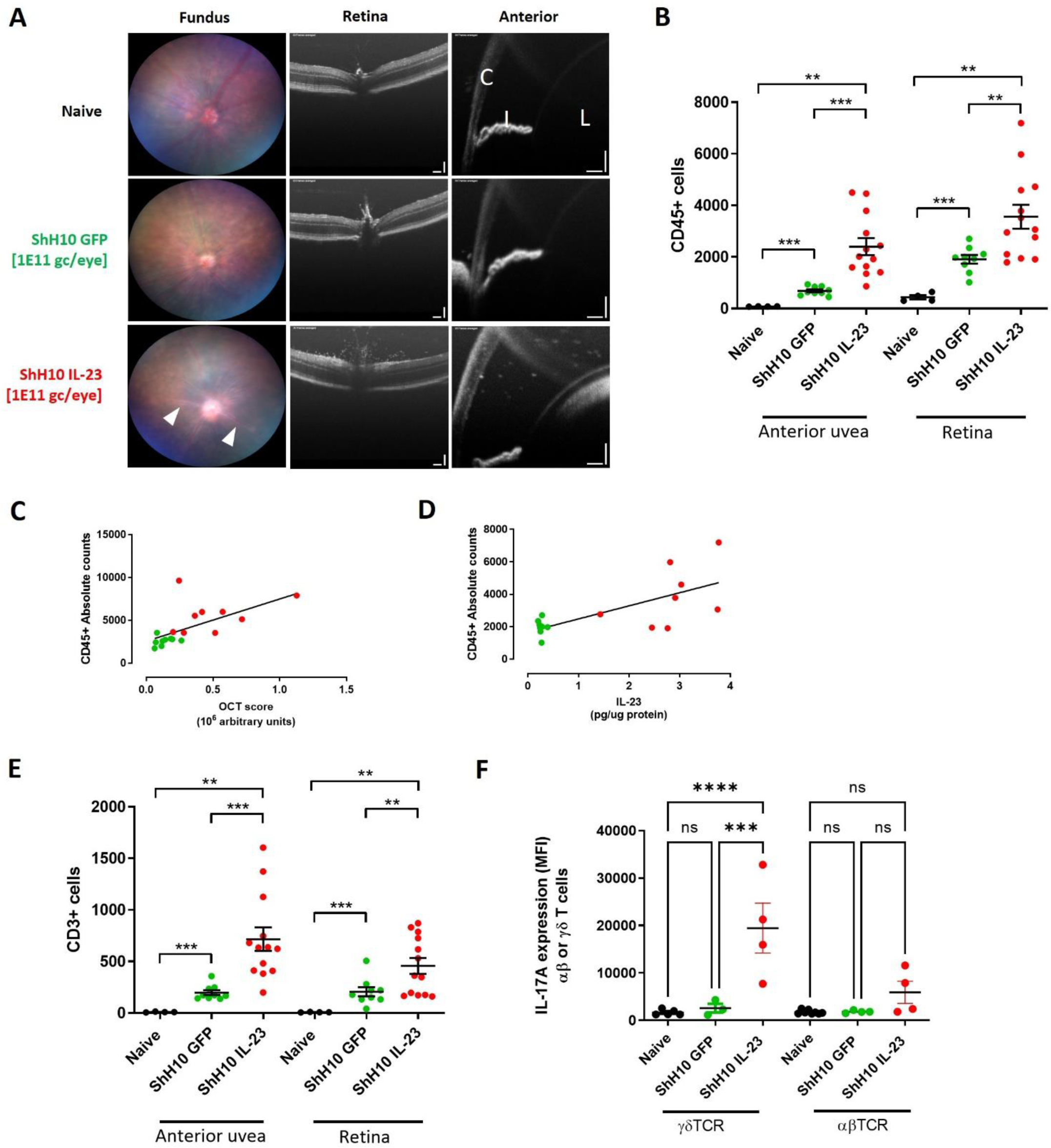
Ocular IL-23 expression drives increased CD45+ infiltration in the AU & retina. Wild-type C57BL/6J mice received intravitreal injection of 1×10^11^ vector genomes of ShH10_IL23, with ShH10_GFP (control AAV) administered to the contralateral eye, and clinically monitored until day 12. Enucleated eyes dissected and anterior uvea and retina samples prepared for flow cytometric immune phenotyping. **(A)** Representative fundus and OCT images of the retina and anterior uvea. White arrow heads indicate peri-vascular sheathing (fundus), posterior (vitreous) and anterior (aqueous) cell infiltrate, respectively in eyes receiving ShH10_IL23 virus. * - optic nerve, C – cornea, I -iris, L – lens; OCT scale bars: 100μm. **(B)** Absolute CD45+ cell counts on day 12 from anterior uvea and retina samples from naïve (n=4), ShH10_GFP (n=9) & ShH10_IL23 (n=13). Absolute counts correlated with **(C)** OCT clinical disease score and **(D)** IL-23 cytokine (protein) expression (n=9/. **(E)** Total live CD3+ counts in anterior uvea and retina on day 12. **(F)** Intracellular IL-17A expression (MFI – mean fluorescence intensity) following ex vivo restimulation from γδTCR+ or αβ TCR+ T cell subsets in response to ShH10_IL-23 (n=4 eye from naive, ShH10_GFP or ShH10_IL-23 injected eyes). Statistical analysis; One-way ANOVA; Data expressed as means +/- SEM; ns = not significant, ** = *P*<0.01, *** = *P*<0.001, **** *P*<0.0001. Data shown is combined from 2 independent experiments **(B & E)** or from single representative experiment **(C, D & F)**.

In eyes receiving ShH10_IL23 virus, increased CD45+ counts correlate with higher clinical disease severity **(Figure 3C)** and expression levels of IL-23 protein detected by ELISA from tissue supernatants **(Figure 3D)**. Phenotypic analysis of the CD45+ population shows that adaptive CD3+ cells (both CD4+ and CD8+) are the predominant cell type recruited to the eye in response to IL-23 expression **(Figure 3E**; see also **Supplementary Figure 2)**. *Ex vivo* restimulation (PMA/Ionomycin) of anterior uvea drives significantly higher expression of IL-17A from the γδ T cell subset compared to αβ T cell subset in eyes receiving ShH10_IL-23 **(Figure 3F)**. Whilst αβ T cells are the predominant IL-17A producers in the anterior uvea, no differences were observed between the different conditions (naïve, ShH10_GFP, or ShH10_IL-23). This indicates that whilst the absolute number of IL-17+ γδ T cells are not statistically increased in response to local IL-23 expression, the resident cells are activated and produce more IL-17A. Furthermore, the number of IFNγ−producing αβ T cells is increased with control or ShH10_IL-23 virus, and no change is detected in the γδ T cell compartment **(Supplementary Figure 3A-D)**.

To evaluate long-term outcome of IL-23 overexpression in the eye, we also monitored mice over an extended time-course, until day 50 post-intravitreal injection. Clinical imaging (fundus and OCT) demonstrates that eyes receiving the ShH10_IL-23 progress to develop a chronic and persistent inflammation, in contrast to control (ShH10_GFP) eyes that remain normal. FACS of combined AU and retina confirms disease severity, with a 10-fold increase in the total CD45+ and CD3+ absolute counts compared to the day 12 cell numbers **(Supplementary Figure 4)**.

### CD3+ γδTCR+IL-23R+ are necessary and sufficient to drive IL-23 mediated inflammation

These data support that tissue-resident T cells respond to IL-23 in the local ocular environment to elicit inflammation (uveitis). To test the dependence of uveitis on T cells we went on to identify whether Th cells (Th17 or T γδ17) or other cell types such as ILC3 were the central drivers for the early response to localized IL-23 expression. All these cell lineages express functional IL-23 receptors [22, 27, 53] but unlike Th17 or γδ17 T cells, ILC3 do not require Rag2 expression for their development [64].

C57BL/6J and B6.Cg-Thy1 (Rag2 knock-out) mice, receiving 1×10^11^vg ShH10_IL-23 in one eye and control ShH10_GFP vector in the contralateral eye were clinically monitored until day 12 and cellular infiltrate in the anterior and posterior compartments assessed by flow cytometry. In Rag2 deficient animals, administration of the ShH10_IL-23 (or control ShH10_GFP) did not replicate any of the clinical inflammatory changes (perivascular sheathing, vitreous infiltrate) that are observed with IL-23 overexpression in wild-type C57BL/6J mice **(Figure 4A)**. The absence of clinical inflammation in the Rag2 mice is confirmed by flow cytometry analysis, demonstrating no recruitment of CD45+ infiltrating cells to either ocular compartment **(Figure 4B)**. This data suggests that T cells (Th17 or T γδ17) are the key effector cell type responding to IL-23 in the eye.

**Figure 4:**
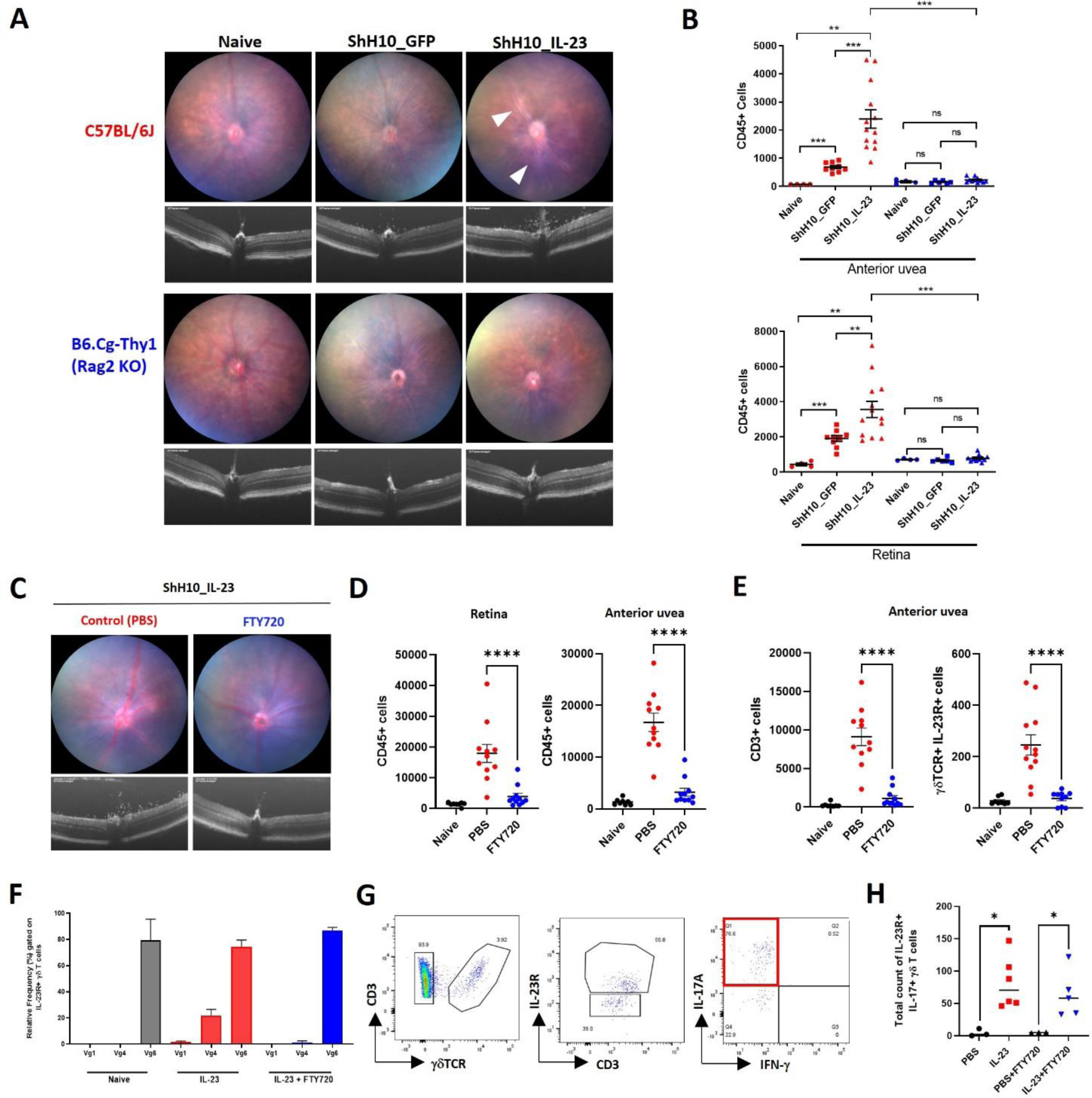
CD3+ γδTCR+IL-23R+ are both necessary and sufficient to drive IL-23 mediated inflammation. C57BL/6J and B6.Cg-Thy1 (Rag2 knock-out) received intravitreal injection of 1×10^11^ vector genomes of ShH10_IL23 or ShH10_GFP vectors. Clinical monitoring (fundus and OCT) performed until day 12. Eyes were enucleated and prepared for flow cytometry to quantify immune cell infiltration in the anterior uvea and retina. **(A)** Representative clinical images show perivascular sheathing (white arrow heads) and vitreous infiltrate present in C57BL/6J eyes receiving ShH10_IL23 only. **(B)** Total live CD45+ cell counts (anterior uvea or retina) from C57BL/6J or Rag2 KO mice at day 12 (naïve (n=4); ShH10_GFP (n=6-9) or ShH10_IL-23 (n=6-13); data combined from 2 independent experiments). IL-23R-eGFP(+/-) mice were bilaterally injected with ShH10_IL23 (1×10^11^ vector genomes) and allocated to groups (n=6) for repeated oral dosing with Fingolimod (FTY720; 10mg/kg) or vehicle control, administered on alternate days following AAV injection. On day 12 eyes were evaluated using clinical imaging and enucleated for flow cytometric analysis of cell infiltrate. **(C)** Representative images show inflammation absent in eyes receiving FTY720 treatment. **(D, E)** Total live CD45+ counts in the anterior uvea and retina, and CD3+ and γδTCR+IL-23R+ cells in the anterior uvea. Statistical analysis, One-way ANOVA; Data expressed as means +/- SEM; ns = not significant, ** = *P*<0.01 and *** = *P*<0.001. **(F)** Relative frequency of γδTCR+ expressing Vγ1, Vγ4 or Vγ6 in naïve and ShH10_IL23 injected eyes receiving vehicle or FTY720 treatment. **(G, H)** Following *ex vivo* restimulation, representative gating of anterior uvea γδTCR+IL-23R+ cells in response to ShH10_IL23 to show relative frequency of IL-17A expressing cells is equivalent between the FTY720 and vehicle treatment groups. Statistical analysis, Mann-Whitney test; * = *P*<0.05.

Recognizing the primed effector phenotype and pathogenic capacity of the resident IL-23R+ γδ+ T cells in healthy tissue, we next explored how local IL-23 expression influences effector function (IL-17A production) and the recruitment of peripheral immune cells. IL-23R-eGFP reporter mice injected with ShH10_IL23 or vehicle (PBS) in the contralateral eye, were treated with a repeated oral dosing regimen of FTY720 [10mg/kg], a potent Sphingosine-1-phosphate receptor 1 (S1PR1) agonist that hinders T cell migration to inflammatory sites including the eye [65]. At day 12 post AAV injection, no clinical disease or CD45+ infiltrate was evident confirming that FTY720 treatment inhibits the recruitment of peripheral immune cells to both the retina and anterior uvea in response to local IL-23 expression **(Figure 4C&D)**. Analysis of the anterior uvea highlights that as FTY720 inhibitory action prevents the increase in total CD3+ number, the resident CD3+IL-23R+ γδ+ T cell population size remains comparable to the naïve tissue. In contrast, absence of S1PR1 blockade leads to a reciprocal increase in the number of CD3+IL-23R+ γδ + cells **(Figure 4E)**, as a mixed population of cells comprising Vγ1, Vγ4 and Vγ6 subsets **(Figure 4F)**. Following *ex vivo* stimulation, IFN-γ expression was not detected **(Figure 4G)**, with the frequency of IL-23R+ γδ+IL-17A+ cells similar between the vehicle and FTY720 treatment groups **(Figure 4H)**.

### Human anterior uvea contains CD3+ γδTCR+ cells capable of producing IL-17A

Our data from the mouse demonstrates that the anterior uvea contains resident CD3+IL-23R+ γδ+IL-17A+ T cells, that when activated drive the recruitment of peripheral CD45+ immune cell populations and promote inflammation in both ocular compartments. To understand whether a similar resident population could be identified in man, we next characterised the immune cell biology in the equivalent insertional regions of the human eye using post-mortem tissue obtained from healthy donors. Confocal microscopy reveals CD3+ T cells are present in key structural areas of the anterior uvea (AU) including the iris, ciliary body, iridocorneal angle (ICA) and the limbal sclera as far as the peripheral cornea **(Supplementary Figure 5)**.

To determine whether these T cells were in the eye tissue and not intravascular artefact, eye tissue sections were co-stained with CD34 and podoplanin (PDPN), markers of vascular and lymphatic endothelium, respectively. This confirmed that tissue-resident T cells are located within the folds of the ciliary processes **(Figure 5A)**; ciliary body proximal to the ciliary muscles **(Figure 5B)**; ciliary region of the iris, proximal to the anterior epithelium and dilator muscles of the posterior border **(Figure 5C)**; the border of the ICA, arranged alongside PDPN+ trabeculae cells **(Figure 5D)**, and in the cribriform layer of the TM adjacent to wall of Schlemm’s canal **(Figure 5E)**. No T cells were observed in the uveal meshwork or corneoscleral meshwork except where trabeculae merge with the iris root and penetrate the ciliary body at the ICA.

**Figure 5:**
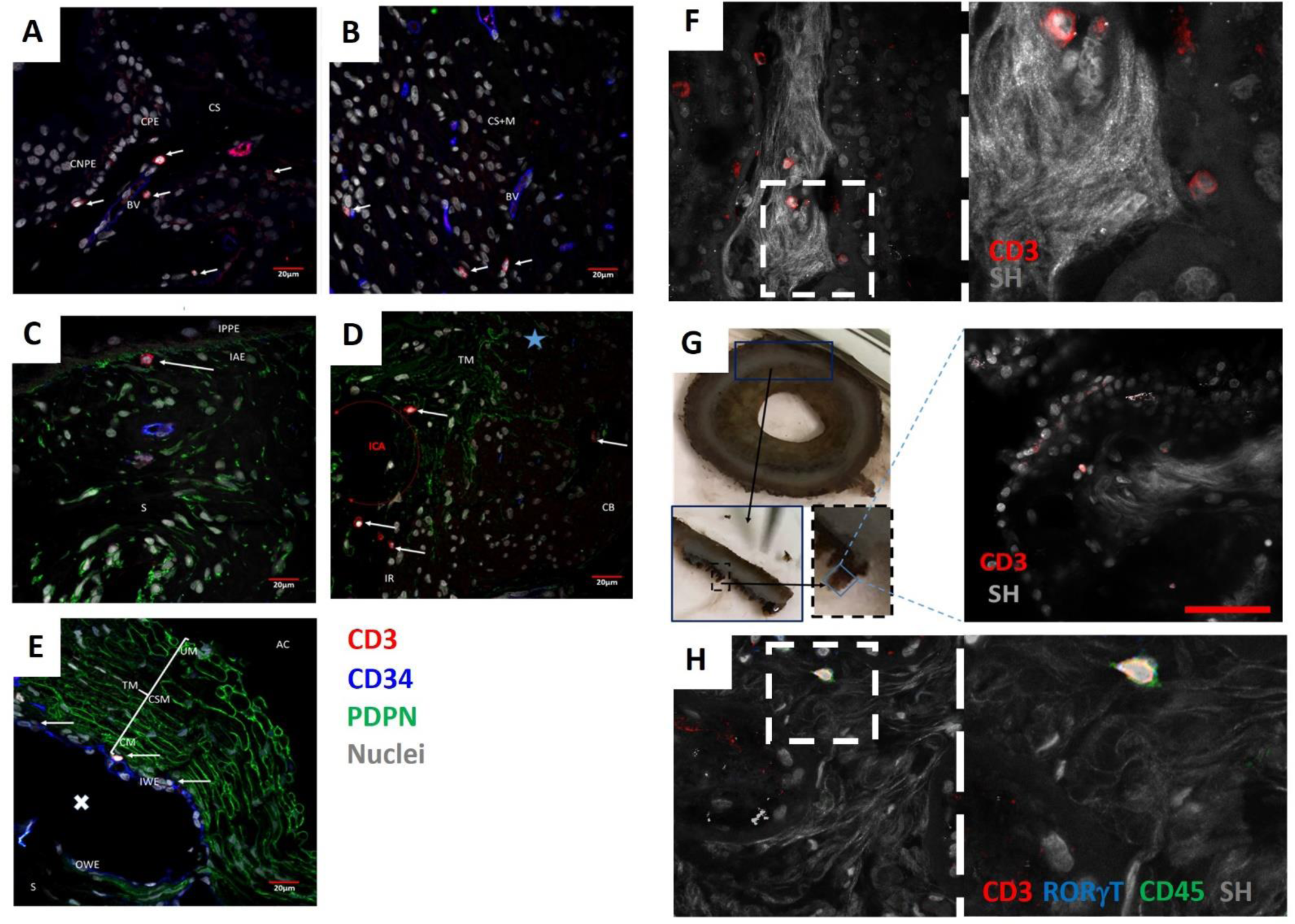
The human anterior uvea contains tissue resident CD3+ cells. FFP tissue sections prepared from an enucleated human eye with a healthy anterior chamber were co-stained with antibodies to CD3 (red), CD34 (blue), podoplanin (green) and Hoechst nuclear stain (grey). Representative confocal immunofluorescence (IF) images showing extravascular location of CD3+ T cells in the **(A)** ciliary processes; **(B)** ciliary body proximal to the longitudinal ciliary muscles; **(C)** within the ciliary region of the iris proximal to the anterior epithelium and dilator muscles of the posterior border layer; **(D)** at the border of the ICA and arranged alongside PDPN+ trabeculae cells; **(E)** in the cribriform layer of the trabecular meshwork adjacent to the inner wall of Schlemm’s canal. AC – anterior chamber, CNPE – ciliary non-pigmented epithelium, CPE – ciliary pigmented epithelium, CS – ciliary stroma, BV – blood vessel. Red scale bar indicates 20μm. **(F)** Anterior tissue section stained for CD3 (red) and Collagen type 4 (Col IV; green), with higher magnification images in red squares, showing ciliary body and limbal scleral regions, respectively. **(G)** Representative immunofluorescence image showing merged expression of second harmonic resonance (SHR) collagen fibres and nuclei (grey), and CD3 (red) within a major ciliary process. Blue hashed square highlights magnified region. Photographs detailing dissection of the major ciliary process from healthy human anterior uvea for whole mount staining, with SHR image showing merged expression of Col IV and CD3. Red scale bar indicates 50μm. **(H)** Images to show merged expression of SHR collagen fibres, CD3, and RORγT (blue). Blue hashed square highlights magnified region. All SHR images captured at x40 magnification.

To assess the relationship of CD3+ cells with the collagenous vascular core of the ciliary processes, and the basement membrane of the ciliary pigmented epithelium (CPE), high magnification multiphoton imaging was utilised to generate second harmonic resonance images of the collagen fibres from coronal sections of healthy tissue **(Figure 5F)**. This demonstrates T cells are closely associated with the fibres that constitute the dense collagen core of the ciliary processes. Most T cells were observed at the centre of the collagenous cores, but cells are also located proximal to fine collagen fibres, inserting into the CPE. This was validated by whole mount imaging of ciliary body tissue to negate the potential for displacement of T cells during the sectioning process **(Figure 5G)**. Despite low frequency, second harmonic imaging also enabled us to identify RORγT+ expression of CD3+ cells within the normal ciliary body **(Figure 5H)**.

To exclude the potential artefact of peripheral blood contamination, we compared the relative frequencies of T cell subsets present in post-mortem anterior uvea (AU) and scleral (S) tissue, with peripheral blood mononuclear cells (PBMCs) isolated from normal healthy controls **(Figure 6A)**. In comparison to blood, AU and S samples contain a significantly lower proportion of CD4+, with a higher frequency of CD8+ **(Figure 6B)**. Flow cytometry highlights AU and S tissues possess a diverse array of leukocyte subsets, including CD3+CD4-CD8-T cells **(Figure 6C)** Whilst the relative frequency of this population remains unchanged compared to PBMCs, a small subset of CD3+ T cells were γδTCR+ **(Figure 6D)**. Extended immune-phenotyping of the CD3+ γδTCR+ population compared to PBMCs revealed a higher frequency of AU CD3+ T cells expressing CCR6 and CD161 **(Figure 6E)**, producing IL-17A following ex vivo stimulation **(Figure 6F)**.

**Figure 6:**
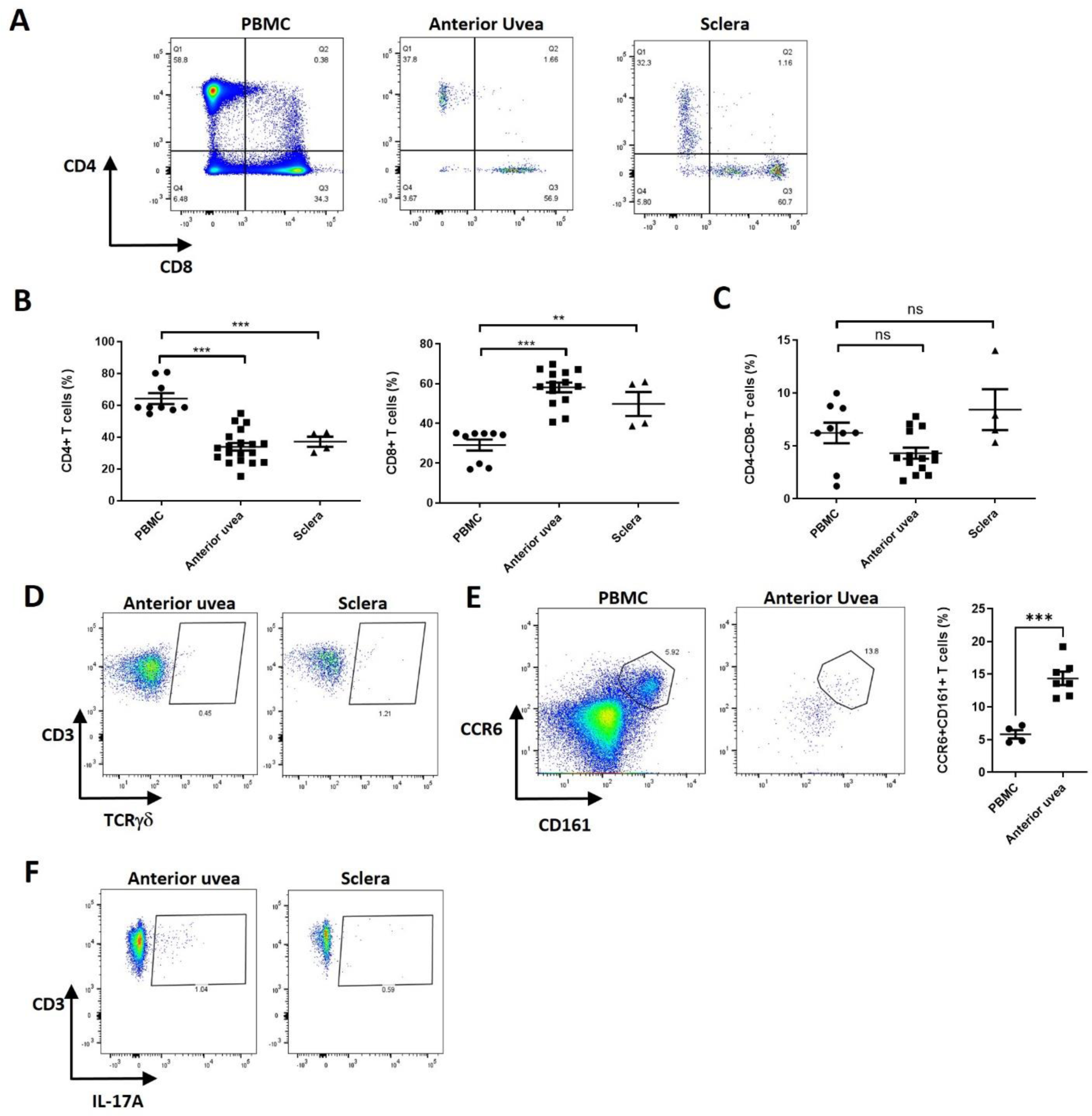
CD3+γδTCR+ cells in the anterior uvea and sclera can produce IL-17A. Flow cytometric analysis was used to phenotype T cell subsets from post-mortem anterior uvea (AU) and sclera (S) tissue samples and compared with peripheral blood mononuclear cells (PBMCs) isolated from normal healthy controls. **(A, B)** Representative FACS plots showing CD3+ populations positive for CD4+ or CD8+ isolated from AU, S and PBMCs and compiled frequency data for the percentage of different CD3+ T subsets. **(C, D)** Relative frequency of CD3+CD4-CD8-cells observed in ocular tissues (AU & S) and PBMCs, and representative FACS plots to show CD3+γδTCR+ cells in AU and S. Samples: anterior uvea (n = 19; data obtained from 7 independent experiments), sclera (n = 4; data obtained from 2 independent experiments), and control blood (n = 9; data obtained from 4 independent experiments). Mean percentage is represented by the central line with error bars showing SEM ** = P<0.01 and *** = P<0.001 (unpaired Student’s two-tailed t-test). **(E)** Representative FACS plots to demonstrate gating of CCR6+ CD161+ cells in PBMC and AU samples, and graph showing relative frequency of this population within the CD3+ T cell pool. Samples: PBMC (n=4) AU (n=7) *** = P>0.001 (unpaired Student’s two-tailed t-test. **(F)** FACS plots of AU and S following *in vitro* stimulation with PMA/Ionomycin to demonstrate a fraction CD3+ cells express IL-17A.

Collectively, this human data reveals that the healthy anterior tissues contain populations of T lymphocytes which are located within collagenous cores at the centre of the ciliary processes, within the ciliary body and the sclera.

## Discussion

Here we present new evidence that a previously unidentified resident population of IL-23R+ γδ T cell, primed to rapidly respond to IL-23 exists within the anterior uvea in mice and humans. Consistent with previous reports of the functional presence of CD3+CD4-CD8-γδTCR+IL-23R+ T cells at musculoskeletal entheseal tissues and aortic root [21, 26], CD3+CD4-CD8-γδTCR+IL-23R+ T cells also constitute the majority of resident anterior compartment IL-23R+ T cells. Despite being present at a low frequency in normal healthy tissue, these cells exhibit a “primed” phenotype with an intrinsic capacity to secrete IL-17A as pathogenic effectors. *In vivo*, localized induced ocular expression of the IL-23 cytokine demonstrates resident CD3+ γδTCR+IL-23R+ cells are both required and sufficient following activation to drive the recruitment of peripheral CD45+ infiltrating cells and promote inflammation (uveitis) in the mouse eye. Furthermore, data from study of human anterior uvea tissue, demonstrates extravascular location of resident CD3+γδTCR+ cells with capacity to generate IL-17A, identifying potential IL-23R mediated responsiveness in man.

The healthy uvea (anterior and posterior regions) of the eye have traditionally been considered devoid of lymphocytes, outside circulating cells in blood vessels [66], with occasional studies reporting the occurrence of conventional αβ TCR+ T cells in the normal iris and ciliary body [67, 68]. Understanding of tissue-resident lymphocyte populations has expanded over the past few years, and resident memory T-cells (T_RM_) as well as other cell types including innate lymphoid cells (ILCs) and non-classical T cells expressing unique TCR heterodimers (γδ T-cells) have been identified [69, 70]. Typically, high numbers of γδ T cells are found at barrier surfaces, including the conjunctiva (lining of the eyelid and globe) which possesses a resident population involved with regulating mucosal immunity and barrier homeostasis in host defense [71]. Recent reports using a transgenic TCR reporter mouse suggested the presence of γδ T-cells in tissues proximal to the limbal sclera (without any reference to vasculature) [26], and present within the choroidal vasculature of healthy human donor eyes [72]. Taken together, this supports our data that T cells are present in the uvea.

Using non-pigmented (albino) and IL-23R reporter transgenic naïve mice, we demonstrate the anatomical location and phenotype of this novel resident T cell population present in the healthy anterior uvea and adjacent tissue. Imaging the whole intact anterior compartment revealed clusters of CD3+ cells located within the peripheral cornea, sclera, and ciliary body. Immunofluorescence on tissue sections from IL-23R-eGFP mice pinpoints their extravascular niche by demonstrating that cells reside within the pars plicata of the ciliary body, proximal to longitudinal ciliary muscle and within the folds proximal to the ciliary epithelium, the iris, and stromal layer of the adjacent sclera. The ciliary body, as a circular muscle positioned immediately behind the iris, functions to produce ocular aqueous fluid but is also connected via zonular fibres enabling changes of crystalline lens shape (accommodation). Furthermore, this region of the anterior compartment is also proximal to extra-ocular muscle insertions. Accordingly, these intra- and extra-ocular tissues are intimately involved with mechanical movement and stress and are analogous to the MSK entheses: we therefore consider them as “ocular entheses”.

To enhance deeper-immune-phenotyping of this rare cell population within the naive eye we utilized multi-parameter/spectral flow cytometry techniques. Enzymatic digestion to liberate immune cells, revealed the healthy mouse anterior uvea contains a relatively small population of CD3+CD4-CD8-γδTCR+IL-23R+ T cells. Importantly, the γδTCR+IL-23R+ population exhibits a “primed” phenotype, confirmed through surface expression of (CD44^high^CD62L-CD27-CCR6+), and intracellular expression of RORγT, which drives the secretion of pro-inflammatory IL-17A upon stimulation. These cells share characteristics of pathogenic effectors, including the same steady-state phenotype as the γδ T cell populations described at the entheseal regions of the axial skeleton and aortic root in the mouse [21, 26].

γδ T cells are a unique T cell subpopulation, rare in secondary lymphoid organs but enriched in many peripheral tissues with an intrinsic capacity to express large amounts of effector cytokines (IFN-γ or IL-17A) that shape local immune responses [73]. Different waves of γδ T cell progenitor subsets, classified based on their somatic T cell receptor gene rearrangements (TCRγ; Vγ chain) and functional potential are produced during specific developmental windows in the thymus before selectively homing to different organs [48, 74–76]. Whilst the extent of their functional roles is still being defined, homing to peripheral sites is widely considered to provide a mechanism that expands spatial immune responsiveness in tissues not served by conventional αβ T cells. In addition to immune surveillance, recent studies highlight other roles in steady-state physiology with evidence of involvement in neuronal synaptic plasticity in the CNS [77].

The γδTCR+IL-23R+ population identified in the normal mouse anterior tissues display Vγ6 TCR homogeneity (no expression of Vγ1 or Vγ4), combined with their ‘primed’ surface phenotype and cytokine profile, indicating these cells represent a long-lived IL-17-producing γδ T cell population similar to the skin, entheseal tissues, lung, reproductive tract and brain [49, 50, 76]. Elaboration of the embryonic development window and timing when these cells are seeded in the anterior eye tissue, will help inform as to whether this resident population exert other physiological roles, akin to CNS γδ T cells.

The current characterisation has only been undertaken in adult mice (6-8wks), therefore examining the extent of their resident tissue longevity, alongside the influence of age-associated decline in their functional immune phenotype in the anterior uvea will be important. Ageing leads to substantial compositional changes in the peripheral lymph node γδ T cell pool in mice, skewed toward an expansion and polarisation of cells toward IL-17 producing Vγ6 γδ T cells alongside age-related increases in tumour incidence [78]. In humans, an altered γδ T cell usage and increased effector phenotype is observed with age [79]. If similar is true for tissue-resident populations in the anterior compartment and tissues associated with AxSpA, such age-related changes may explain the link to increased risk for extra-articular manifestations in AxSpA including AAU.

To understand how resident γδTCR+IL-23R+ cells could act as pathogenic effectors in the context of AAU, we employed IL-23 overexpression, previously achieved through systemic DNA mini-circle injection [21, 26, 27, 80]. Development of the ShH10-IL-23 AAV vector for intravitreal delivery facilitated ocular restricted expression of IL-23 to interrogate the role of the resident population in driving inflammation. Inflammation elicited in the mouse eye replicates many clinical features that correlate to human disease, namely posterior (retina) inflammation, and cellular infiltration in both the vitreous and aqueous compartments. Following injection, constitutive secretion of IL-23 led to rapid onset of inflammation, followed by a chronic and persistent disease phenotype not observed or attributed to ShH10_GFP response. Use of AAV to model, probe and evaluate therapeutics for human inflammatory ocular disease is an expanding area [81, 82], and in the current context provides a tool to interrogate pathogenicity of IL-23 alone in a disease that is accompanied by elevated concentrations of serum and AqH IL-23 [39, 41, 42, 83].

Taking an iterative approach to highlight the effector function of the rare population of T cells we deployed Rag2 deficiency and S1PR1 antagonism (Fingolimod treatment), that unequivocally demonstrated the resident CD3+ γδTCR+IL-23R+ population is both necessary and sufficient to drive uveitis. FACS demonstrates IL-23 expression correlates to the level of immune cell infiltrate, with CD45+ populations comprising elevated numbers of CD3+ T cells (both CD4+ & CD8+), and recruitment of monocytes, macrophages, dendritic cells, B cells and neutrophils.

Anterior eye tissue contained significantly higher numbers of CD3+ γδTCR+ T cells, despite the relative frequency of cells exhibiting the pre-activated effector phenotype in response to IL-23 remaining unchanged when compared to either control AAV injected or naïve eyes. Our interpretation was that the resident population lacks proliferative capacity in response to IL-23 stimulation, leading us to consider whether the expanded number was due to peripheral recruitment. Extended phenotyping of CD3+ γδTCR+IL-23R+ cells support this hypothesis, revealing a heterogeneous γδ T cell pool comprising Vγ1, Vγ4 and Vγ6 subsets. Furthermore, evidence which corroborates their recruitment following activation, is shown in analysis of anterior tissues taken from the mice receiving fingolimod intervention (pharmacological-induced lymphopenia), where a normal frequency (equivalent to naïve) and homogeneity Vγ6+ γδ subset is maintained. We speculate that the most likely origin of the Vγ1 & Vγ4 cells recruited to the eye are from local peripheral lymphoid organs, as these tissues are the recognised seeding location of these subsets during the post-natal wave of immune development [76]. Further approaches to delineate the γδ T cell phenotype and composition within the local draining lymph nodes, including transcriptomic and cell tracking studies would be informative.

Nonetheless, these cells do not secrete IFN-γ following *ex vivo* stimulation, emphasizing their functional capacity at this location as γδ IL-17-producing (γδ17) T cells that can orchestrate and promote tissue inflammation. Whilst γδ17 T cells are considered exclusively derived from foetal thymus, studies indicate they undergo homeostatic proliferation and self-renewal in peripheral tissues to maintain their numbers [74, 78, 84, 85]. Furthermore, under specific conditions including TCR stimulation and the presence of IL-1β and IL-23 cytokines, Vγ4+ γδ17 T cells also have the potential to develop and expand in the adult mouse [86].

The observations of IL-23R mediated responsiveness of a previously unknown resident γδ T cell population in the mouse, raises the exciting potential that a similar population also exists in humans. In this study, we provide early exploratory evidence that CD3+γδTCR+ cells with capacity to secrete pro-inflammatory IL-17A exist at equivalent insertional regions of the human eye. Tissue sections highlight CD3+ cells located throughout the anterior uvea, including the iris, trabecular meshwork (TM), sclera and the ciliary body, intimately associated with collagen-rich core of the ciliary processes.

Despite the low detectable frequency on tissue sections examined, second harmonic tissue imaging identified CD3+RORγT+ co-expression. However, recognising the challenges associated with tracking IL-23R and RORγt expression in human cells, our FACS phenotyping instead evaluated CCR6 and CD161 expression on CD3+ cells obtained from anterior tissues [87]. Further work to attribute IL-23R responsiveness and provide comprehensive phenotype of resident T cell populations within the human anterior compartment is required, but single-cell multi-omics datasets also provide evidence of lymphocyte populations within the ciliary body [88, 89]. In the future, approaches harnessing transcriptomic and enhanced immunophenotyping will provide clarity on whether the putative IL-23R+ γδ T cells observed in mouse also exist in humans. In support, emerging evidence from single-cell RNA-sequencing of aqueous biopsies taken from uveitis patients reveals that infiltrates in HLA-B27-associated uveitis are dominated by unconventional T cells (γδ T cells) and myeloid cells [90].

In summary, using murine models we have shown that anatomical sites that are typically inflamed in AAU are primed to rapidly react to IL-23 by the presence of this previously unidentified population of IL-23R+ resident cell. *In vivo* exposure to IL-23 is sufficient to induce highly specific ocular inflammation in the absence of Th17 cells and with rapid kinetics. We propose that the *ocular enthesis* is therefore a functional IL-23–responsive anatomic site, similar to the MSK entheseal tissues, gut and lung, which also contain innate IL-23R expressing cells [91], primed to respond immediately to IL-23. In the context of AxSpA related AAU, promising results using IL-17 antagonists (Secukinumab & Ixekizumab) or combined IL-12/23 biologics (Ustekinumab) suggests modulating these pathways in patients requiring long-term control can provide avenues to intervene for both disease manifestations [92, 93].

## Methods

### Sex as a biological variable

Equal numbers of sex-matched mice were used both for characterization of naive tissue and in disease (involving ShH10_IL23 administration) experiments.

### Animals

Adult C57BL/6J were purchased from Charles River Laboratories, UK. B6(Cg)-Tyrc-2J/J (Albino), B6.Cg-Thy1 (recombination activation gene (Rag)2 KO) and IL-23R-eGFP mice were supplied from established breeding colonies at the University of Bristol, UK. Homozygous IL-23R-eGFP^GFP/GFP^ mice [53] were mated with C57BL/6J mice to generate heterozygous reporter mice for experiments. All mice were used at 6-12 weeks of age. All mouse strains were confirmed as negative for the Rd8 mutation [95], and were housed under specific pathogen free conditions with food and water *ad libitum*.

### Study Approval

All procedures were conducted in concordance with the United Kingdom Home Office licence (PPL PP9783504) and were approved by the University of Bristol Ethical Review Group. The study also complied with the Association for Research in Vision and Ophthalmology (ARVO) Statement for Use of Animals in Ophthalmic and Vision research.

### Synthesis of mouse ‘hyper-IL-23’ cytokine sequence

The sequence synthesis was performed by GenScript, Piscataway, USA. Mouse IL-23 artificial hyperkine (a term coined by the DNAX Research institute that describes two protein subunits connected by a synthetic linker - GSGSSRGGSGSGGSGGGGSKL), consisting of mouse IL-12p40 and p19 as one sequence [60], with flanking restriction enzyme sites and ligated into a pD10-CMV plasmid. In animal models, this ‘hyper-IL-23’ cytokine sequence has been shown to drive pathology strongly resembling human autoimmune inflammatory conditions including psoriasis and AS [19, 21]. pCMV-eGFP, was also used in this study, and was a gift from Connie Cepko (Addgene plasmid #11153; http://n2t.net/addgene:11153; RRID: Addgene_11153).

### AAV Production

AAV vectors were either purchased from Vector Biolabs (PA, USA) or manufactured at the University of Bristol as previously described [96]. In brief, recombinant ShH10 serotype particles were produced through triple-plasmid transfection using PEI transfection reagent into 293T-HEK cells. ShH10 particles were bound to a 1-mL HiTrap AVB Sepharose column (GE Healthcare, USA) and eluted with 50 mM glycine (pH 2.7) into 1 M Tris (pH 8.8). Vectors were desalted and concentrated in PBS-MK to a concentration of 1 x 10^13^ genome copies per milliliter (gc/mL) using a Vivaspin 4 (10 kDa) concentrator. Vector genome titers were determined by quantitative real-time PCR using probes binding to ITR sequences **(Supplementary Table 1)**. An amplicon-based standard series of known concentration was used for sample interpolation. Preparations were certified as endotoxin <5 EU/mL by Pyrotell-T kinetic turbidimetric endotoxin test (Associates of Cape Cod, MA, USA). Vectors used in the study were termed ShH10_IL-23 (expressing ‘hyper-IL-23’ cytokine) or ShH10_EGFP (control vector expressing GFP).

### Intravitreal injection, in vivo imaging and treatment interventions

Prior to any procedure, pupils were dilated using topical tropicamide 1% w/v and phenylephrine 2.5% w/v (Minims; Chauvin Pharmaceuticals, Romford, UK), before anaesthesia with 2% isofluorane (Piramal Critical Care, West Drayton, UK). AAV was administered by intravitreal injection (2 uL/eye), using a 33G needle on a microsyringe under direct visualization (Hamilton Company, Reno, NV, USA). Retina fundal imaging and optical coherence tomography (OCT) scans were captured using Micron IV (Phoenix Research Laboratories). Clinical scoring using vitreous cell densitometry was performed on circular retinal OCT images centred on the disc using ImageJ software [97].

For FTY20 treatments, mice were orally dosed with FTY720 (Fingolimod-hydrochloride; Caymen Chemicals, Ann Arbor, Michigan) 10mg/kg or vehicle control, administered on alternate days following AAV injection. Mice were assigned to treatment groups (FTY720 or vehicle) in a constrained randomised order within blocks, dependent on cage allocations and litter they were derived from.

### Immunostaining, flow cytometry and ImageStream analysis

#### Murine ocular tissue

##### Dissection and dissociation

Following enucleation, single eyes were dissected in 100µL of ice-cold PBS. Using a limbal incision the posterior segment was removed, whole retina and vitreous extracted and together with the dissecting fluid (PBS) transferred into a 1.5mL Eppendorf tube. The tissue was mechanically dissociated by rapping the tube across an 80-well standard rack 10 times.

For anterior tissue, following lens removal, the iris, ciliary body and limbal sclera was first mechanically dissociated in a culture dish using scissors, before enzymatic digestion in 0.5mL DMEM containing 5 mg/mL type II collagenase (Worthington, # LS004202) and 0.2 mg/mL DNase I (Roche, #11284932001) for 45 minutes at 37°C undergoing constant, gentle agitation. The enzymatic digest was stopped by adding 0.5mL of complete media (DMEM containing 10% FCS), centrifuged at 250g for x 10mins and cell pellets resuspended in 100ul of cold PBS.

Retina and anterior cell suspensions were transferred into a 96-well 60-μm cell filter plate (Merck Millipore) and washed with 150µL of PBS. The plate was centrifuged at 1200 rpm for 5 min, the supernatant aspirated and cells resuspended in 0.1% bovine serum albumin (BSA) fluorescence-activated cell sorting buffer and transferred into a 96-well V-bottom plate for immunostaining.

#### Flow cytometry

##### Cell surface marker staining

Cells were incubated with purified rat anti-mouse CD16/32 Fc block (1:50; 553142, [2.4G2], BD) for 10 min at 4°C before incubation with fluorochrome-conjugated monoclonal antibodies against mouse cell surface markers **(Supplementary Table 2)** at 4°C for 20 min. Cells were washed and resuspended in 7-aminoactinomycin D (Thermo Fisher Scientific) for dead cell exclusion.

##### Intracellular cytokine staining

Anterior uvea or retinal cell suspensions were initially stimulated for 2 hours with 50 ng/ml phorbol myristate acetate (PMA) and 500 ng/ml Ionomycin, and then for a further 2 hours with the addition of 1 mg/ml Brefeldin and Monensin (BD Biosciences) in complete medium. Cells were fixed, permeabilized with the Cytofix/perm solution (BD Biosciences) and stained with intracellular cytokine antibodies **(Supplementary Table 3)** LIVE/DEAD fixable cell staining kit (Thermofisher Scientific, USA) was used to exclude dead cells from analysis.

##### Cell acquisition

Cell suspensions were acquired using a fixed and stable flow rate for 2.5 min on a four-laser Fortessa X20 flow cytometer (BD Cytometry Systems). Compensation was performed using OneComp eBeads (01-1111-41, Thermo Fisher Scientific). Seven two-fold serial dilutions of a known concentration of splenocytes were similarly acquired to construct a standard curve and calculate absolute cell numbers [98]. Analysis was performed using FlowJo software (Treestar).

##### ImageStream Analysis

Following surface receptor staining, anterior uvea cell suspensions were fixed, permeabilized using the FOXP3 staining kit (Ref: 00-5523, eBioscience) and stained with RORγT (1:40; 12-6981-82, [B20] eBioscience). Cells were washed with PBS containing 2% FCS twice. From naive eyes, x4 anterior uvea cell suspensions were pooled, and image-based flow cytometry analysis conducted in Oxford using the ImageStream MkX. DAPI (2µg/mL; BD Pharmingen; 564907) was added to samples immediately prior to acquisition. 10,000 events per sample were collected and analysed using the IDEAS software.

#### ELISA

A mouse IL-23 Quantikine ELISA kit (Biotechne, USA) was used to quantify levels of IL-23 according to manufacturer’s instructions. Homogenised ocular samples were spun at 13,000 rpm for 10 minutes and the supernatant typically diluted between 1:10 to 1:1000 prior to testing. Samples were analysed with technical triplicates unless limited material precluded this. This assay has a detection range of 15.6 - 1,000 pg/mL, recognizing natural and recombinant mouse IL-23, but not free p19 or p40 subunits.

#### Human ocular tissue

Human donor eye material surplus to corneal transplantation (without recorded ocular disease) was obtained from National Health Service (NHS) Blood and Transplant Services after research ethics committee approval (REC#: 07-H0706-81), with experiments conducted according to the Declaration of Helsinki and in compliance with UK law. Partial globes were supplied (transported and processed) within 36hrs from time of death. Patients with known inflammatory or infectious condition of the entheses were excluded from the study.

##### Dissection and cell dissociation

Briefly, the entire anterior uvea was removed by cutting around the circumference from the choroid to the base of the ciliary body (avoiding any sclera and retinal attachment from the ora serrata. Following this, a ∼2mm cut around the circumference of the limbal sclera was made (which included any muscular attachment sites). Both tissues were washed twice in Dulbecco’s Modified Eagle’s Medium (DMEM; Gibco, #41965-039), cut into small ∼1mm^3^ pieces and enzymatically digested in DMEM containing 5 mg/mL type II collagenase (Worthington, # LS004202) and 0.2 mg/mL DNase I (Roche, #11284932001) for 120 minutes at 37°C undergoing constant, gentle agitation. The digest was then washed with complete media and filtered twice through a 70μm cell strainer (Corning, # CLS431751-50EA). Single cell suspensions were stained with the Zombie NIR™ Fixable Viability KIT (780nm of excitation; Biolegend, #423106) at a 1/250 dilution in PBS (Biolegend, #420201) for 20 minutes at 4°C.

##### Cell surface receptor staining

Cells were washed (Cell staining buffer; Biolegend) before resuspension in Human TruStain FcX™ blocking agent, before incubation with fluorochrome-conjugated monoclonal antibodies against human cell surface markers **(Supplementary Table 4)** for 20 minutes on ice. Cells were washed twice then placed into 400µL cell staining buffer in FACS tubes. Compensation controls were made using UltraComp eBeads (eBioscience). Cells were analysed on a 3 laser (405nm, 488nm, and 640nm) BD LSR Fortessa X-20 flow cytometer (BD Biosciences). Medium or high flow rates were used. FACSdiva and FlowJo software (BD Biosciences) were used for analysis of data. Compensation was performed using OneComp eBeads (eBioscience, San Diego, USA, 01-1111-42).

##### Intracellular cytokine staining

Intracellular cytokine staining of T-cells was performed by stimulating cell suspensions for 2 hours with 50 ng/ml phorbol myristate acetate (PMA) and 500 ng/ml Ionomycin then for a further 2 hours with the addition of 1 mg/ml Brefeldin and Monensin (all from Sigma Aldrich, Dorset, UK) in D10 medium. Following surface antigen staining, the Transcription Factor Buffer Set (BD Biosciences, Oxford, UK) was used according to the manufacturer’s protocol for intracellular staining. A specific dilution of each primary conjugated antibody was used **(Supplementary Table 5)**. Following this, cells were washed twice then placed into 200µL 0.1% w/v BSA in PBS in FACS tubes.

### Histology

#### Murine ocular tissues

##### Immunofluorescence imaging on tissue sections

For immunofluorescent imaging, mouse eyes (B6(Cg)-Tyrc-2J/J (Albino), C57BL/6J or IL-23R-eGFP(+/-) were fixed with 4% paraformaldehyde, frozen in optical cutting temperature compound (VWR, PA, USA), and sectioned at 12-μm intervals. Slides were incubated with a 1:1,000 dilution of DAPI (Sigma Aldrich, UK) and mounted in fluorescence mounting media (Agilent Technologies, CA, USA) before imaging on an EVOS FL microscope (Thermo Fisher Scientific, UK) or Leica SP5 Confocal microscope (Leica Microsystems, Germany).

##### Whole tissue anterior segment Lightsheet imaging

Following euthanasia, 8-week-old *B6(Cg)-Tyrc-2J/J* mice underwent cardiac perfusion with 4% Paraformaldehyde. Eyes were enucleated and the anterior segment isolated by dissection along the ocular equator. The anterior segments were then placed overnight in 1:4 dilution of BD Cytofix/Cytoperm in PBS at 4°C. The following day this was washed once in PBS then incubated for 24 hours at room temperature in 1x BD Wash Perm buffer with 5% normal goat Serum (Jackson Immunolabs). This was then replaced with 1x BD Wash Perm buffer, with diluted AlexaFluor 647 conjugated rat anti-mouse CD3 antibody (1:100; 101010; Clone [17A2]) and incubated for 3 days at room temperature. The tissue was washed for 4 hours then placed into Ce3D clearing medium [47] overnight, before mounting in Ce3D medium and imaging on a Z1 Lightsheet system (Zeiss) acquiring the pre-set 633nm and 488nm channels. Images underwent 3D reconstruction using Imaris version 8.0 (Oxford Instruments).

#### Human ocular tissues

Human ocular tissues were fixed in 10% buffered formalin, before paraffin embedding, sectioning at 4 μm thickness onto adhesive glass slides (Leica, # 3800050) and baked at 60°C for 30 mins and then at 37°C for a further 60 mins.

##### Antigen retrieval

Slides were incubated at 60°C for 60 min, and tissue sections subjected to deparaffinization and target retrieval steps (heat-mediated antigen retrieval at pH 6) using an automated PT Link instrument (Dako). Antibody staining was performed using the EnVision FLEX visualization system with an Autostainer Link 48 (Dako). Antibody binding was visualized using FLEX 3,3′-diaminobenzidine (DAB) substrate working solution and haematoxylin counterstain (Dako, UK). Primary antibodies against human CD3 (1:100; #A045229-2) and CD68 (1:400; #M0876) were obtained from Dako, UK. Images were acquired using an inverted microscope (Axiovision software -Zeiss). Due to extremely small sample size of the specimens, only four images were acquired: two at x20 and two at x40 magnification.

##### Immunofluorescence imaging on tissue sections

This protocol was adapted from a previous paper [99]. Briefly, following antigen retrieval, tissues were blocked in 5% normal goat serum (Sigma-Aldrich) in PBS for 60 min in a humid chamber at room temperature. After removal of blocking solution, sections were incubated with the primary antibody cocktail diluted in 5% normal goat serum in PBS for 2 hours at room temperature. Details of primary antibodies used for immunofluorescence are listed in **(Supplementary Table 6)**. Sections were washed (3x PBS–Tween 20 (PBST) for 5 min), the incubated with secondary antibodies each diluted 1:200 in 5% normal equine serum (Sigma-Aldrich) in PBS for 2 hours. The secondary antibodies were Alexa Fluor goat anti-mouse IgG2a or IgG2b or goat anti-rabbit IgG (Life Technologies) and goat anti-mouse IgG1 (Southern Biotech). Sections counterstained with 2 μM POPO-1 nuclear counterstain (Life Technologies), diluted in PBS containing 0.05% saponin (Sigma-Aldrich) for 20 min. Tissue autofluorescence was quenched with a solution of 0.1% Sudan Black B (Applichem) in 70% ethanol for 10 min. Slides were mounted using fluorescent mounting medium (VectaShield). For negative controls, the primary antibody was substituted for universal isotype control antibodies: cocktail of mouse IgG1, IgG2a, IgG2b, IgG3, and IgM (Dako) and rabbit immunoglobulin fraction of serum from non-immunized rabbits (Dako).

### Second Harmonic Imaging

Multi-photon images were acquired on a Zeiss LSM 710 laser-scanning confocal coupled to an inverted Zeiss Axio Observer.Z1 microscope. A 40x/1.3 NA oil immersion objective was used with immersion media RI: 1.518. Two-photon excitation was performed using a Chameleon Vision II, Ti:Sapphire laser (Coherent, 680 - 1080 nm, pulse width: 140 fs at peak, repetition rate: 80MHz). Second Harmonic Generation (SHG) imaging was performed by setting the two-photon at 760 nm to reach an excitation wavelength of 380 nm in the sample. Images were acquired with non-descanned detectors (NDD) using a filter set equipped with a long pass dichroic LP 445 nm and a bandpass filter (BP380-430).

### Statistical analysis

Statistical tests applied and n sample numbers are detailed in the figure legends. Data were analysed using GraphPad Prism^®^ software (v9.4). Variance was compared using an F-test. Data with equal variances were analysed using the unpaired Students two-tailed t-test. Non-parametric data (e.g. ocular cell infiltrate) were analysed using one-way ANOVA and Tukey’s multiple comparison test, with data expressed as means +/- SEM; ns = not significant, ** = *P*<0.01, *** = *P*<0.001, **** *P*<0.0001.

## Supporting information

Combine supplementary figures 1-5

## Data availability

All relevant information about data is available directly from the corresponding author. Values for all data points in graphs are reported in the Supporting Data Values file.

## Acknowledgements

This work was supported by funding from the Underwood Trust, Sight Research UK, NIHR Biomedical Research Centre at University of Oxford, NIHR Biomedical Research Centre at Moorfields Eye Hospital and UCL Institute of Ophthalmology, and Johnson & Johnson. CJC is funded as a Wellcome Clinical Research Career Development Fellow (224586/Z/21/Z). We thank Alison Young & John Pooley for ShH10 AAV preparations (Bristol), Joanne MacDonald for assistance in collecting control blood samples (Oxford), Nadia Halidi for second harmonic resonance imaging (CRG) and Julia Lewis for providing the 17D1 hybridoma (Yale). We thank Lindsay Nicholson and Gareth Jones (Bristol) for critical reading of the manuscript. We also acknowledge the flow cytometry and imaging facilities based at the University of Bristol, Kennedy Institute and Sir William Dunn School of Pathology at Oxford. All members of the ORBIT Consortium <u>ORBIT — The Kennedy Institute of Rheumatology (ox.ac.uk)</u>

## Author contributions

JS & ADD conceived the study. RH, AW, CJC, SGD designed, performed, and or analyzed experiments. SEC provided samples of consented human tissue. DAC, JS, ADD, SK, PCT designed and or analyzed experiments and supervised research. RH, CB, JS, ADD & DAC wrote the manuscript, with contributions from all other authors. RH & AW contributed equally to this work and are co–first authors.

## Appendix 1

**Supplementary Table 1:**
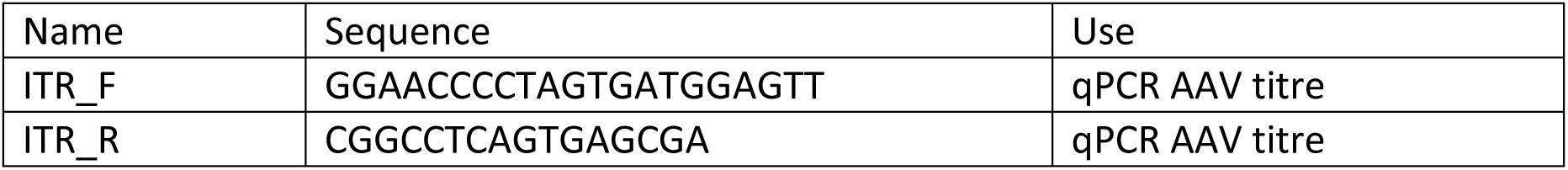
ITR oligonucleotides sequences.

**Supplementary Table 2:** Mouse surface staining antibody panel and dilution.

| Antigen | Fluorochrome | Clone | Manufacturer | Dilution |
| --- | --- | --- | --- | --- |
| CD11c | af488 | HL3 | BD | 1/200 |
| F4/80 | PE | T45-2342 | BD | 1/200 |
| CD4 | PE-Dazzle | GK1.5 | BIOLEGEND | 1/100 |
| CD45 | PE-CY7 | 30-F11 | BD | 1/1500 |
| CD11b | AF647 | M1/70 | BD | 1/100 |
| Ly6C | AF700 | AL-21 | BD | 1/200 |
| CD8 | APC-CY7 | 53-6.7 | BIOLEGEND | 1/20 |
| Ly6G | BV421 | 1A8 | BD | 1/100 |
| CD19 | BV510 | 6D5 | BIOLEGEND | 1/20 |
| CD3 | BV786 | 145-2C11 | BIOLEGEND | 1/40 |

**Supplementary Table 3:**
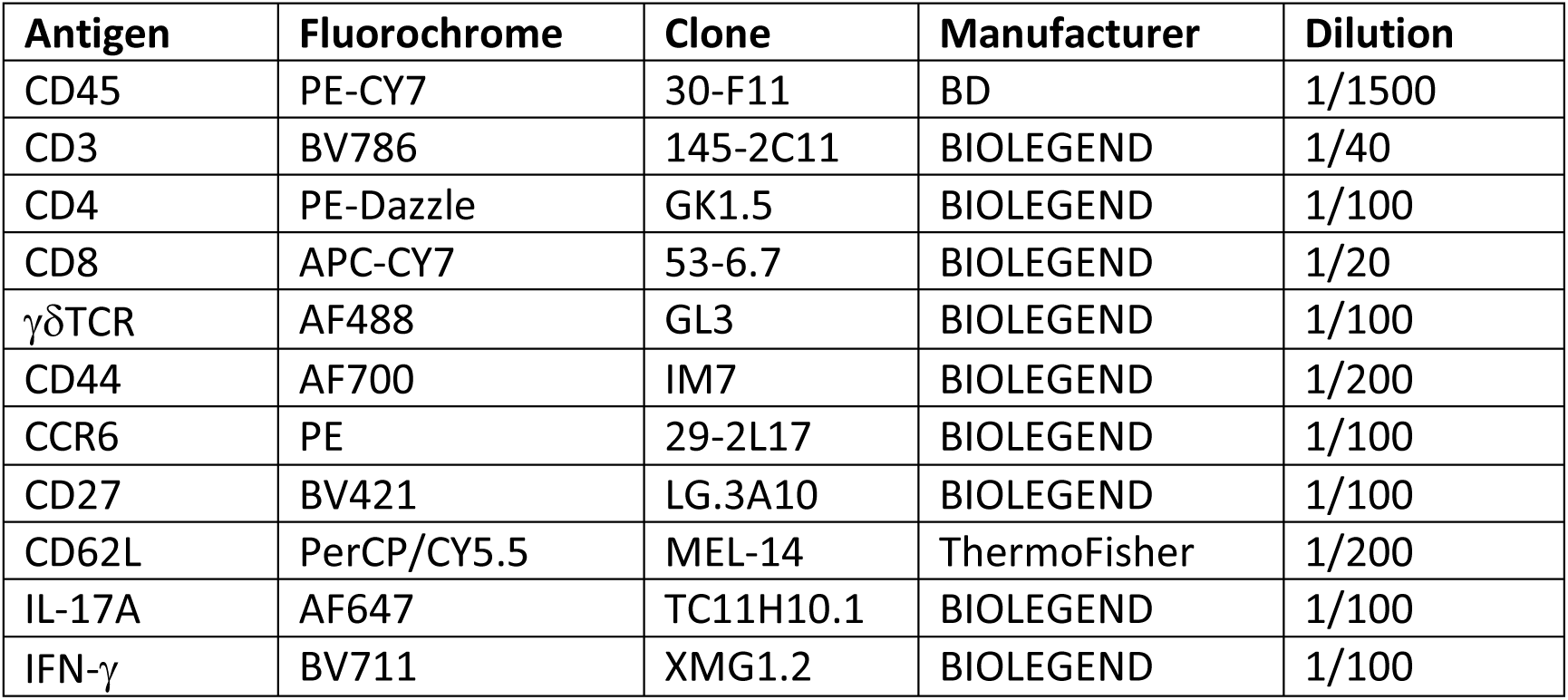
Mouse intracellular flow cytometry panel.

**Supplementary Table 4:** Human surface staining antibody panel.

| Antigen | Fluorophore | Clone | Manufacturer | Dilution |
| --- | --- | --- | --- | --- |
| PANEL 1 - LEUKOCYTES |  |  |  |  |
| CD45 | BV605 | H130 | BIOLEGEND | 1/20 |
| CD3 | AF647 | HIT3a | BIOLEGEND | 1/20 |
| CD14 | FITC | M5E2 | BIOLEGEND | 1/20 |
| CD68 | PE-CY7 | Y1/82A | BIOLEGEND | 1/20 |
| CD11B | BV421 | ICRF44 | BIOLEGEND | 1/20 |
| CD4 | AF700 | RPA-T4 | BIOLEGEND | 1/50 |
| CD8 | PerCP/CY5.5 | RPA-T8 | BIOLEGEND | 1/20 |
| CD20 | PE | 2H7 | BIOLEGEND | 1/20 |
| Viability dye | APC-CY7 | N/A | BIOLEGEND | 1/250 |

| PANEL 2 – TH17 |  |  |  |  |
| --- | --- | --- | --- | --- |
| CD45 | BV605 | H130 | BIOLEGEND | 1/20 |
| CD3 | AF647 | HIT3a | BIOLEGEND | 1/20 |
| CD4 | AF700 | RPA-T4 | BIOLEGEND | 1/50 |
| CD8 | PerCP/CY5.5 | RPA-T8 | BIOLEGEND | 1/20 |
| CCR6 | FITC | G034E3 | BIOLEGEND | 1/20 |
| CD161 | BV711 | DX12 | BD | 1/20 |
| Viability dye | APC-CY7 | N/A | BIOLEGEND | 1/250 |

**Supplementary Table 5:** Human intracellular flow cytometry antibody panel.

| Antigen | Fluorophore | Clone | Manufacture | Dilution |
| --- | --- | --- | --- | --- |
| CD45 | BV605 | H130 | BIOLEGEND | 1/20 |
| CD3 | AF647 | HIT3a | BIOLEGEND | 1/20 |
| CD4 | AF700 | RPA-T4 | BIOLEGEND | 1/50 |
| CD8 | PerCP/CY5.5 | RPA-T8 | BIOLEGEND | 1/20 |
| γδTCR | BV421 | B1 | BD | 1/20 |
| IFN-γ | PE-CY7 | B27 | BD | 1/20 |
| IL-17 | FITC | ebio64DEC17 | THERMO FISHER | 1/20 |
| CD20 | PE | 2H7 | BIOLEGEND | 1/20 |
| Viability dye | APC-CY7 | N/A | BIOLEGEND | 1/250 |

**Supplementary Table 6:** Human tissue immunofluorescence antibody panel.

| Antigen | Clone | Isotype | Host Species | Reactivity | Manufacturer | Dilution |
| --- | --- | --- | --- | --- | --- | --- |
| CD45 | 2B11+PD7/26 | IgG1 | Mouse | Human | Dako | no dilution |
| CD34 | 3C8G12 | IgG2B | Mouse | Human | Proteintech | 1/200 |
| CD3 | POLY | IgG | Rabbit | Human | Dako | 1/200 |
| CD3 | F7.2.38 | IgG1 | Mouse | Human | Dako | 1/200 |
| PDPN | 18H5 | IgG1 | Mouse | Human | Abcam | 1/200 |
| Col T1 | COL-1 | IgG1 | Mouse | Human | Abcam | 1/200 |
| ROR gamma | POLY | IgG | Goat | Human | Abcam | 1/200 |

