## Supplementary material for "IL-23 drives uveitis by acting on a novel population of tissue-resident entheseal T cells": Combine supplementary figures 1-5

**Title**

### Supplementary Figures:

**Supplementary Figure 1: Phenotyping of CD3+ γδTCR+IL-23R+ T cells in the naïve eye**

**Supplementary Figure 2: Infiltrating CD45+ cells comprise both adaptive and innate immune cell types**

**Supplementary Figure 3: αβ T cells are the predominant IL-17 producing cells following ex vivo stimulation**

**Supplementary Figure 4: Extended time-course demonstrates intravitreal delivery ShH10_IL-23 AAV leads to chronic, persistent inflammation at Day 50**

**Supplementary Figure 5: Tissue resident CD3+ T cells present in limbal sclera and ciliary body**

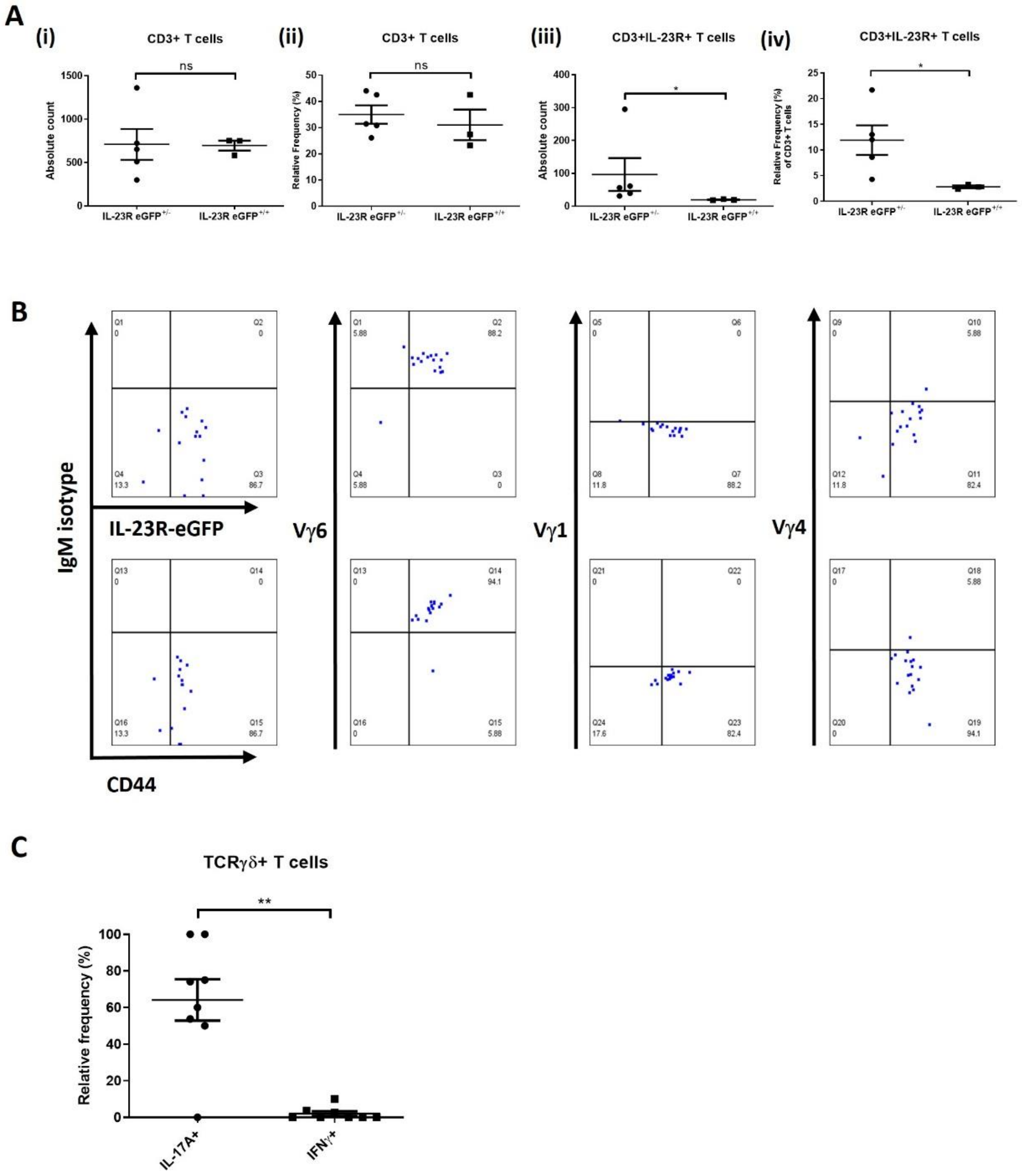


**Supplementary Figure 1: Phenotyping of CD3+ γδTCR+IL-23R+ T cells in the naïve eye.**
Functional IL-23R expression may represent an important determinant in the accumulation and/or residency of ocular CD3+ γδTCR+IL-23R+ T cells. FACS compiled frequency data of pooled left and right anterior tissue collected from IL-23R eGFP+/- (heterozygous) or IL-23R eGFP+/+ (homozygous) mice (n = 5). The absolute count and relative frequency of viable CD3+ T cells ((i) & (ii)) and CD3+γδTCR+IL-23R+ cells. Line represents mean and error bars show SEM. * = *P*<0.05 (unpaired Student's two-tailed t-test). **(B)** Frequencies of anterior CD3+γδTCR+ expressing Vγ1, Vγ4 or Vγ6, as determined by FACS analysis on gated IL-23R+ or CD44+ cells isolated from the naïve anterior uvea of IL-23ReGFP+/- reporter mice. Dot plots depict pooled data (two eyes from single mouse), representative of 2 independent experiments. **(C)** FACS and intracellular cytokine staining of CD3+γδTCR+ cells from anterior uvea samples at Day 12 post-injection of ShH10 IL-23 (1E11vg/eye). Cells were stimulated with PMA/ionomycin and IL-17A and IFN-γ production shown in compiled frequency plots symbols represent pooled left and right eyes from individual mice. Data shown as means ± SEM and are representative of single experiment. ns=not significant, ∗ = P < 0.05 ∗∗ = P < 0.01, unpaired t-test.

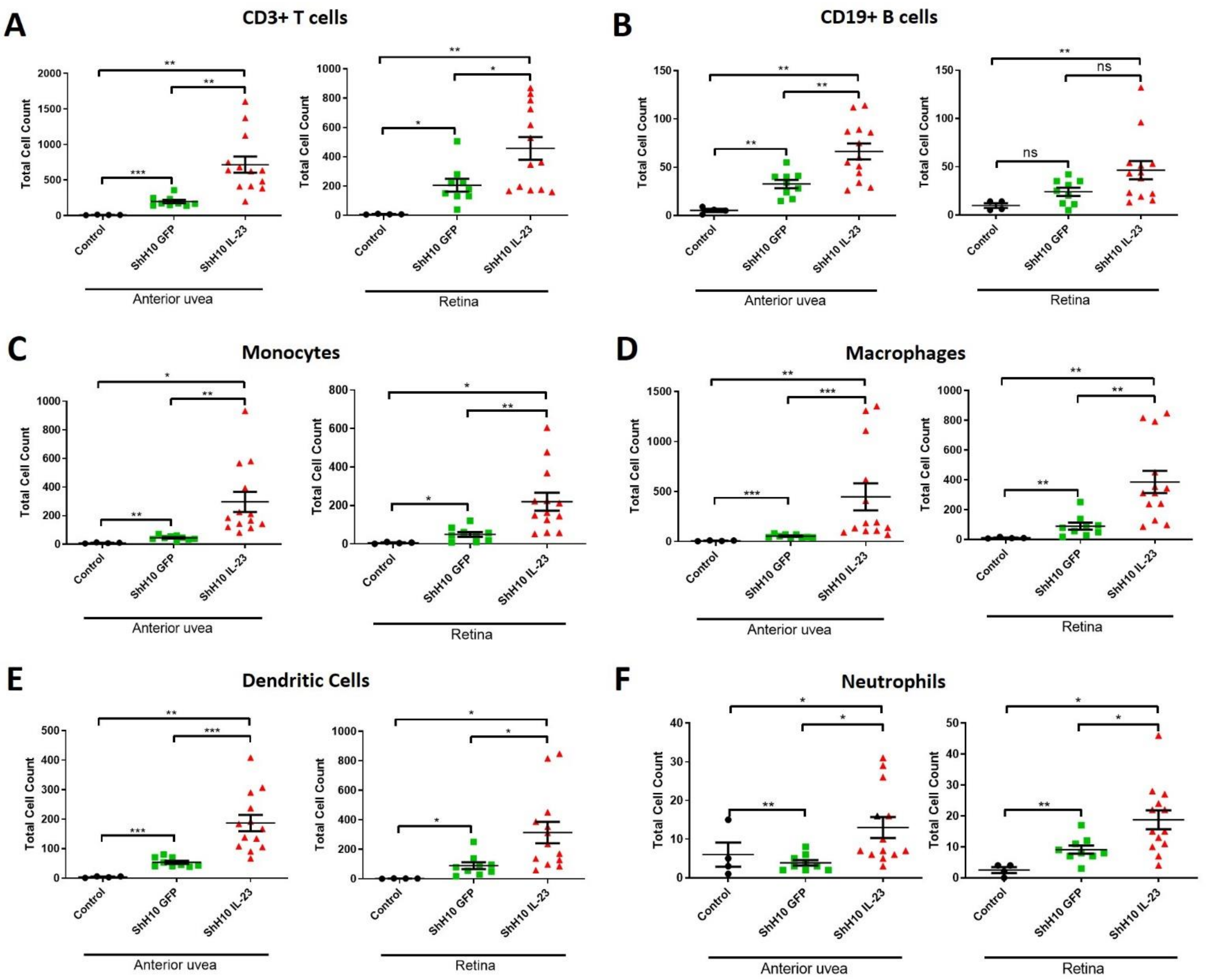


**Supplementary Figure 2: Infiltrating CD45+ cells comprise both adaptive and innate immune cell types.**

Flow cytometric analysis of anterior uvea and retina samples collected at day 12 post intravitreal injection of ShH10.GFP or ShH10.IL-23 [1E11vg/eye]. CD45+ infiltrate comprises both adaptive and innate immune cell populations including **(A)** CD3+ T cells, **(B)** CD19+ B cells, **(C)** CD11b+F4/80- Monocytes, **(D)** CD11b+F4/80+ Macrophages, **(E)** CD11c+F4/80- Dendritic cells, and **(F)** Ly6G+ Neutrophils. Mean count is represented by the central line with error bars showing SEM. * = $P$<0.05, ** = $P$<0.01, and *** = $P$<0.001 (unpaired Student's two-tailed t-test).

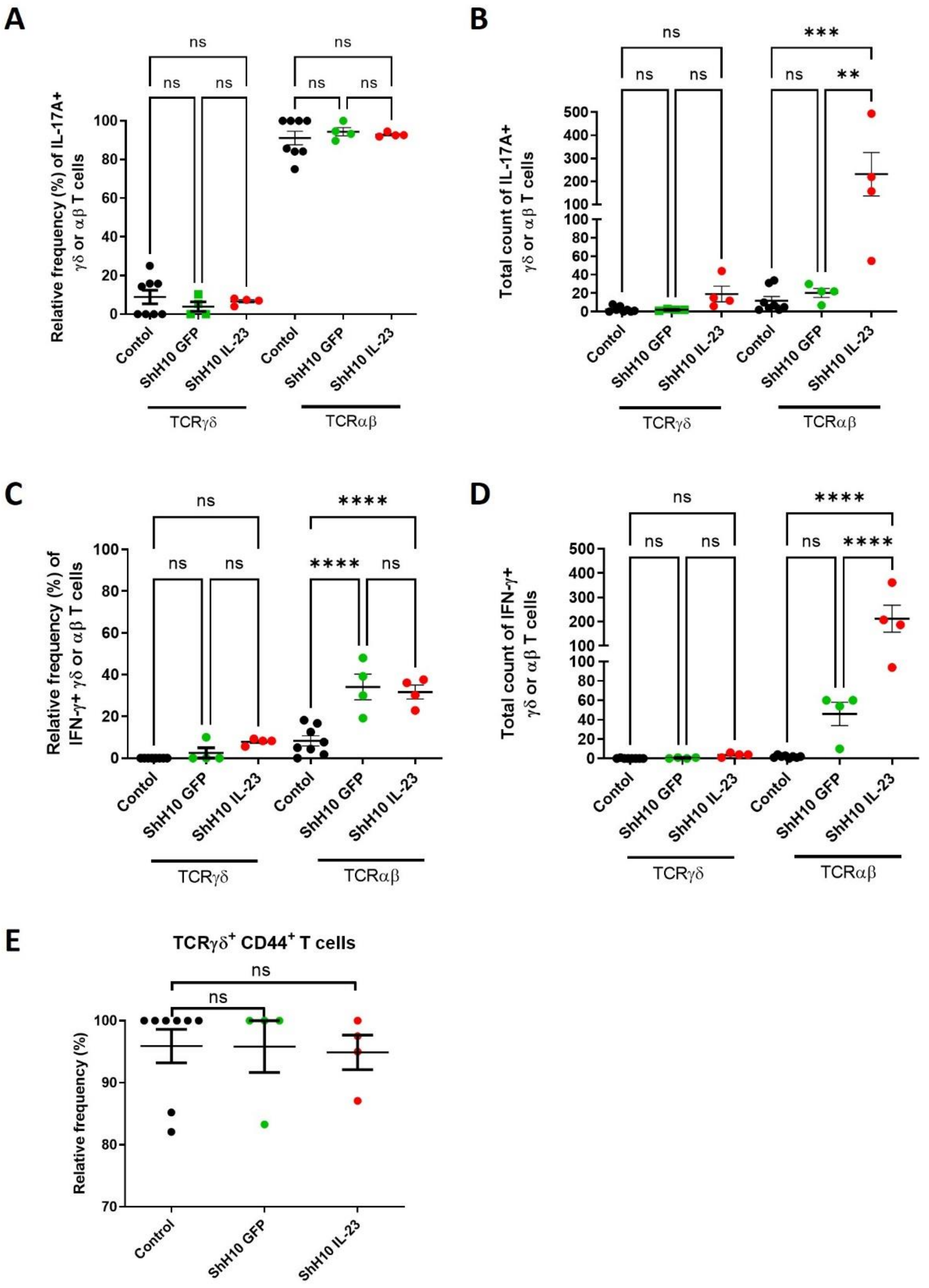


**Supplementary Figure 3: αβ T cells are the predominant IL-17 producing cells following ex vivo stimulation.**
Flow cytometric ICCS phenotyping of αβ and γδ T cell subsets in the anterior uvea. Wild-type C57BL/6J Eyes from naïve controls (un-injected; n=8) or eyes receiving ShH10_GFP or ShH10_IL23 (n=4/group) were dissected at day 12, *in vitro* stimulated (PMA/ionomycin) and stained for surface markers (CD45, CD3, γδ TCR, CD44) and intracellular cytokines (IL-17A and IFN-γ). **(A, B)** Relative frequency and total counts for IL-17A+ αβ and γδ T cell subsets. **(C, D)** Relative frequency and total counts for IFN-γ+ αβ and γδ T cell subsets. **(E)** Relative frequency of CD3+γδTCR+CD27-CD44high cells. Statistical analysis; One-way ANOVA; Data expressed as means +/- SEM; ns = not significant, ** = *P*<0.01, *** = *P*<0.001, **** *P*<0.0001.

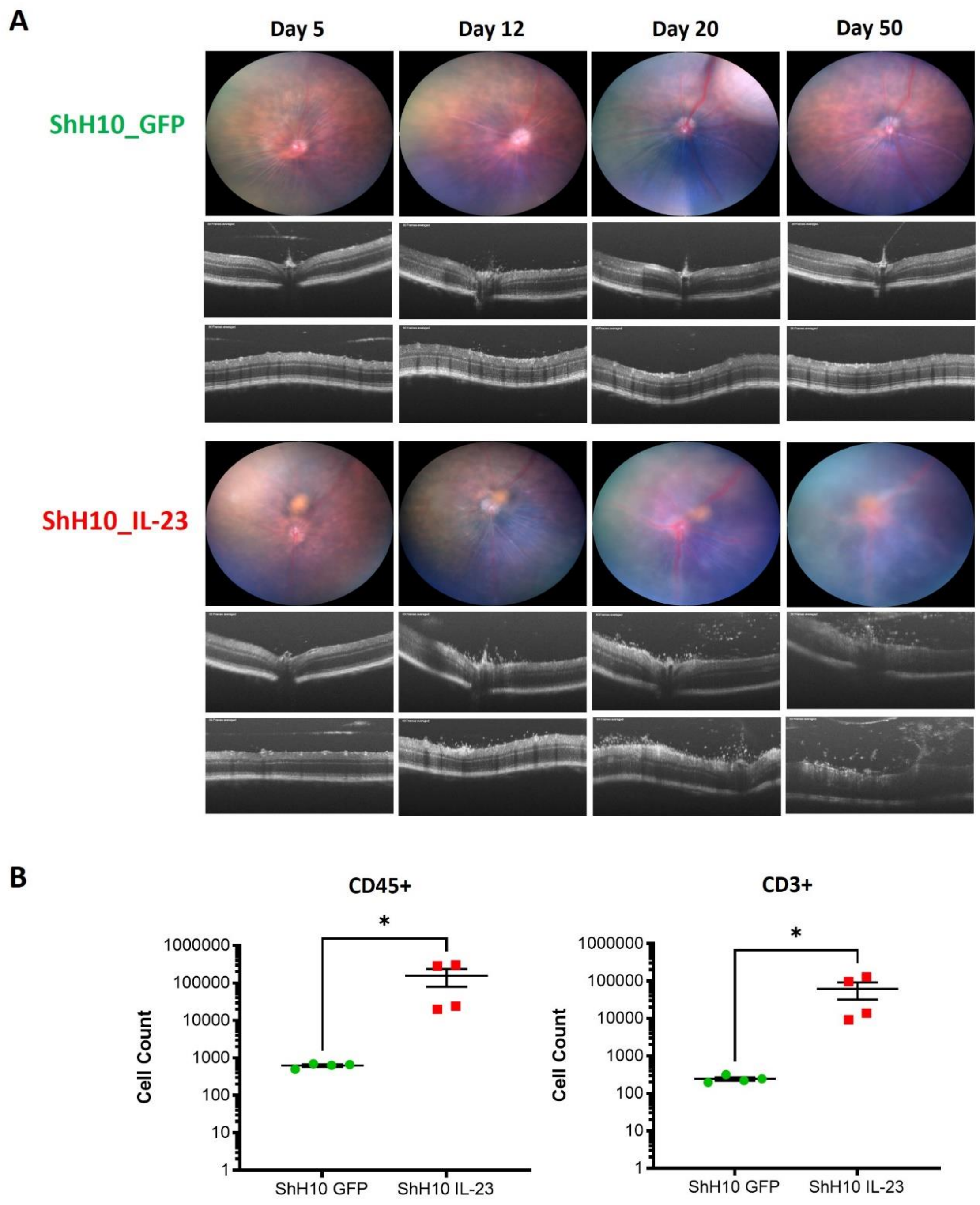


**Supplementary Figure 4: Extended time-course demonstrates intravitreal delivery ShH10_IL-23 AAV leads to chronic, persistent inflammation at Day 50.**

Wild-type C57BL/6J mice (n=4) received intravitreal injection of $1x10^{11}$ vector genomes of ShH10_IL23, with ShH10_GFP (control AAV) administered to the contralateral eye. **(A)** Mice were clinically monitored (Fundus and OCT) and representative images from single mouse are shown for days 5, 12, 20 & 50 post-injection. On day 50, enucleated eyes were dissected and prepared for flow cytometric immune phenotyping. **(B)** Graphs show total live counts for CD45+ and CD3 cells from combined anterior uvea and retina obtained from individual eyes. Statistical analysis: Mann Whitney test; *p<0.05; Data shown as mean ± SEMs and representative of a single experiment.

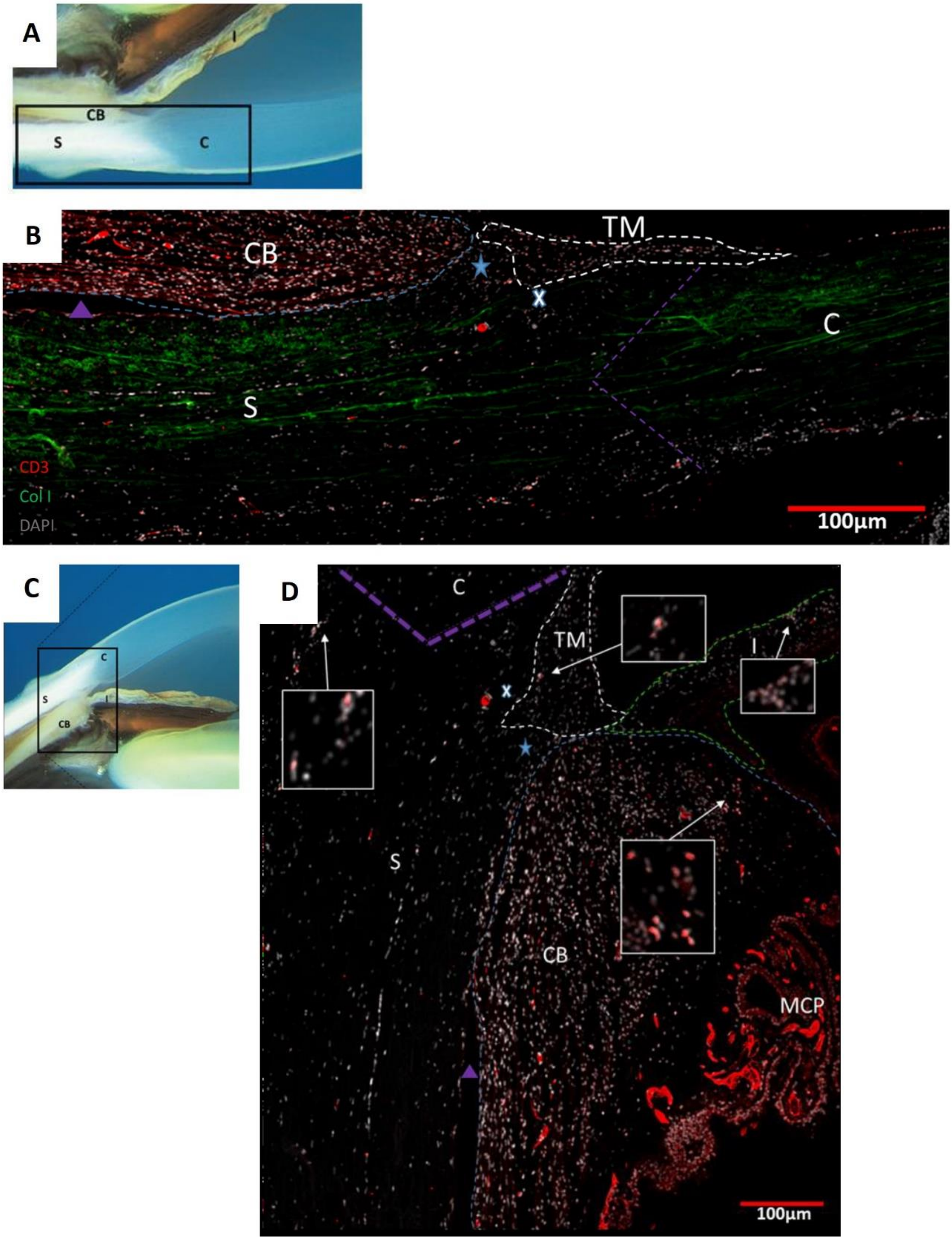


**Supplementary Figure 5: Tissue resident CD3+ T cells present in limbal sclera and ciliary body**

FFPE tissue sections from post-mortem human eyes were obtained from the Liverpool eye bank for immunostaining. **(A, C)** Photographic images to show the tissue location and orientation of the corresponding immunofluorescence image [94]. C – cornea, S – sclera, CB- ciliary body. **(B)** Immunofluorescent staining shows nucleus (grey), collagen type 1 (Col1; green), and CD3+ cells (red). **(D)** Immunofluorescent staining for nucleus (grey) and CD3+ (red) T cells. White arrows and boxes show areas of T cell clusters under digital magnification. Images comprise tile scan composed of 4 x 4 individual images, captured at x10 magnification. C – cornea, S – sclera, purple line – corneoscleral limbus, 'TM' and white line – trabecular meshwork, 'X' – Schlemm's canal, blue star – scleral spur, purple triangle - supraciliary space, 'CB' and blue line – ciliary body, and MCP – major ciliary processes. Red scale bar indicates 100µm.
